# Nanopore direct RNA sequencing detects DUX4-activated repeats and isoforms in human muscle cells

**DOI:** 10.1101/2020.07.27.224147

**Authors:** Satomi Mitsuhashi, So Nakagawa, Martin C Frith, Hiroaki Mitsuhashi

**Affiliations:** Department of Genomic Function and Diversity, Medical Research Institute, Tokyo Medical and Dental University, Tokyo, 113-8510, Japan; Department of Human Genetics, Yokohama City University, Kanagawa, Japan; Micro/Nano Technology Center, Tokai University, Hiratsuka, Kanagawa, 259-1292, Japan; Department of Molecular Life Science, Tokai University School of Medicine, Isehara, Kanagawa, 259-1193, Japan; Artificial Intelligence Research Center, National Institute of Advanced Industrial Science and Technology (AIST), Tokyo, Japan; Graduate School of Frontier Sciences, University of Tokyo, Chiba, Japan; Computational Bio Big-Data Open Innovation Laboratory (CBBD-OIL), AIST, Tokyo, Japan; Department of Applied Biochemistry, School of Engineering, Tokai University, Kanagawa, 259-1207, Japan

**Author notes:** Correspondence to: Hiroaki Mitsuhashi, Department of Applied Biochemistry, School of Engineering, Tokai University, 4-1-1 Kitakaname, Hiratsuka, Kanagawa, 259-1292, Japan.

**Keywords:** DUX4, facioscapulohumeral muscular dystrophy, muscular dystrophy, direct RNA sequencing, long read sequencer, ERV, MaLR

## Abstract

Facioscapulohumeral muscular dystrophy (FSHD) is an inherited muscle disease caused by misexpression of the *DUX4* gene in skeletal muscle. DUX4 is a transcription factor which is normally expressed in the cleavage-stage embryo and regulates gene expression involved in early embryonic development. Recent studies revealed that DUX4 also activates the transcription of repetitive elements such as endogenous retroviruses (ERVs), mammalian apparent LTR-retrotransposons (MaLRs), and pericentromeric satellite repeats (HSATII). DUX4-bound ERV sequences also create alternative promoters for genes or long non-coding RNAs (lncRNAs), producing fusion transcripts. To further understand transcriptional regulation by DUX4, we performed nanopore long-read direct RNA sequencing (dRNA-seq) of human muscle cells induced by DUX4, because long reads show whole isoforms with greater confidence. We successfully detected differential expression of known DUX4-induced genes, and discovered 61 differentially-expressed repeat loci, which are near DUX4-ChIP peaks. We also identified 247 gene-ERV fusion transcripts, of which 216 were not reported previously. In addition, long-read dRNA-seq clearly shows that RNA splicing is a common event in DUX4-activated ERV transcripts. Long-read analysis showed non-LTR transposons including Alu are also transcribed from LTRs. Our findings revealed further complexity of DUX4-induced ERV transcripts. This catalogue of DUX4-activated repetitive elements may provide useful information to elucidate the pathology of FSHD. Also, our results indicate that nanopore dRNA-seq has complementary strengths to conventional short read cDNA sequencing.

## Introduction

The *DUX4* gene, encoding a double homeobox transcriptional factor, was identified in the highly similar 3.3-kb subtelomeric macrosatellite D4Z4 repeats on chromosome 4q35 (1). Since the 4q35 region forms heterochromatin and no transcript was detected, *DUX4* was initially thought to be a pseudogene. However, it was revealed that in FSHD1 patients with D4Z4 repeat contraction, DNA methylation of D4Z4 was reduced and consequently de-repression of DUX4 occurred (2, 3). In FSHD2 patients without D4Z4 contraction, a genetic mutation in *SMCHD1*, which is involved in chromatin modification, reduced D4Z4 methylation and caused DUX4 de-repression (4). Misexpression of DUX4 is thought to be the cause of FSHD because animal models expressing DUX4 demonstrate FSHD-like symptoms (5-8).

Two different DUX4 mRNA isoforms have been identified in human skeletal muscle: a full-length mRNA (DUX4-fl) and a spliced short isoform (DUX4-s) (9). Introduction of DUX4-fl in muscle cells activates a number of cleavage-stage genes and causes cell death (10). In contrast, DUX4-s does not promote transcription and has no cytotoxicity because DUX4-s lacks the C-terminal transactivation domain (11, 12). Consistent with the results in muscle cells, DUX4-fl is transiently expressed in human preimplantation embryos at the four-cell stage and plays a physiological role in regulating gene expression specific to the cleavage stage as a master transcriptional regulator (13).

In addition to gene regulation, DUX4-fl strongly binds to the long terminal repeat (LTR) regions of endogenous retroviruses (ERVs) and mammalian apparent LTR-retrotransposons (MaLRs), and activates transcription of these repetitive elements (10, 14). Since both ERVs and MaLRs have LTRs at their 5’ and 3’ termini, they are categorized as LTR retrotransposons (15). ERVs are thought to have originated from retroviruses, and therefore their genetic structures are similar to those of retroviruses, but MaLR internal sequences are not similar. The LTRs, ranging in size from 300 to 1000 bp, have signals for RNA transcription initiation and 3’-end formation, and sometimes splice donor sites (16). Most ERV and MaLR elements in the human genome are no longer full-length due to deletion, substitution, or mutations over generations. In particular, many single LTRs (so-called “solo LTRs”) are present in the genome, presumably formed by recombination between the flanking LTRs causing loss of the internal region. Previous RNA-seq analyses detected upregulation of ERVs, MaLRs, fusion transcripts of these repetitive elements with a protein-coding gene, and pericentromeric satellite repeats in DUX4-fl-expressing muscle cells (14). Among these repetitive elements, Human Satellite II (HSATII) has been shown to form dsRNA foci and partly play a role in DUX4-fl-induced cytotoxicity (17). In addition, HERVL is activated by DUX4-fl in preimplantation embryos (13), suggesting the activation of repetitive elements by DUX4-fl may have important roles in FSHD pathology and developmental biology.

The findings to date on DUX4-fl-activated repetitive elements are based on data from short read next generation sequencers. However, there may be difficulty in analyzing transcripts comprehensively because there are numerous highly similar sequences in the human genome such as transposable elements (18). Therefore, an alternative approach to complement short-read sequencing may be required for further understanding of the repetitive element-derived transcripts that are activated by DUX4-fl. Long-read sequencing has advantages for analyzing repetitive elements because long enough reads may encompass whole repeat regions, allowing us to specify the genomic location of the repeat (19). In addition, long-read sequencing has better resolution for the transcriptome because it can clearly detect exon connectivity (20, 21). Long read full-length cDNA sequence has been shown to be useful for qualitative and quantitative transcriptome analysis (22, 23). It was also shown that nanopore can sequence native RNA molecules; namely direct RNA sequencing (dRNA-seq) (24). This approach has been also shown to be useful for transcriptome analysis, has more potential power to detect RNA modifications, and avoids PCR or reverse transcription bias (23, 25, 26).

In this study, we sequenced RNAs with polyA tail from DUX4-fl-expressing human muscle cells, using a nanopore direct RNA sequencer. We catalogued DUX4-fl-activated transcripts from repetitive elements with high confidence by statistically analyzing triplicated samples using DESeq2. Nanopore dRNA-seq confirmed the activation of cleavage-stage specific genes in muscle cells. Furthermore, we found novel processed ERV transcripts and LTR fusion transcripts induced by DUX4-fl.

## Results

### Nanopore direct RNA sequencing (dRNA-seq)

We sequenced poly-A enriched RNA extracted from DUX4-fl or DUX4-s overexpressed RD (rhabdomyosarcoma) cells using a single PromethION flowcell per sample in triplicates (DUX4-fl: fl-1, fl-2, fl-3, DUX4-s: s-1, s-2. s-3). Since DUX4-s does not have a transcriptional activation domain and its transcriptional activity is weak (5, 11), we used DUX4-s overexpressing cells as a control. We obtained 5.9 million reads on average, which were mapped to the human reference genome GRCh38 using LAST (https://github.com/mcfrith/last-rna) (Supplementary Table 1) according to the instructions (https://github.com/mcfrith/last-rna/blob/master/scripts.md). Median read length was 824 bases (658-922 bases) and all datasets show similar distribution of read length (Supplementary Figure 1). As with any nanopore sequencing, dRNA-seq is prone to errors. We calculated error probabilities using last-train (27), and aligned the reads to the reference genome using these probabilities. (Supplementary Figure 2). Note that these probabilities contain both actual differences and nanopore errors. We could align approximately 75% of the reads on average to the reference genome, which amounts to 4.6 million reads on average (Supplementary Table 1).

### Differentially expressed genes and repeats

Differentially expressed genes (DEGs) between cells with DUX4-fl and DUX4-s were estimated from nanopore count data, based on a negative binomial generalized linear model and Wald test using DESeq2 (28). dRNA-seq showed remarkable elevation of DUX4-fl induced genes (10, 13), including germline genes or early developmental genes (Figure 1a, b, Supplementary Table 2). This finding is supported by the correlation between mean read count per million (RPM) from our triplicated nanopore data and transcripts per million (TPM) from a public DUX4-fl expressing myoblast dataset (14) (Figure 1c). Gene ontology analysis shows significant enrichment of transcription regulation pathway and cell death pathway (Supplementary Table 3) as reported previously (10). Long reads can show exon connectivity clearly as well as isoform detection. For example, *C12orf50* has several exons and some transcripts show different exon usage (Supplementary Figure 3a). Note that 5’-ends of the dRNA-seq reads do not always agree with the transcription start site (TSS) because nanopore sequencing may end in the middle of the RNA. Nevertheless, we can observe that many reads originate from a novel ERV-derived TSS (arrow) (Supplementary Figure 3b). *C12orf50* encodes an uncharacterized protein highly expressed in testis, which presumably has a role in embryogenesis along with other DUX4-induced genes. *LEUTX* encodes a human embryonic transcription factor, and DUX4-fl induces a functional *LEUTX* isoform with complete homeodomain, but it does not induce another inactive isoform (Supplementary Figure 4) (29). Other isoform-specific transcripts were also detected by dRNA-seq (Supplementary Figure 5). These results show that our nanopore dRNA-seq robustly detects the DUX4-fl inducible gene transcripts. In addition to previously reported DUX4-induced genes, our pipeline identified that expression of TAF11-like macrosatellite array pseudogenes were significantly induced by DUX4-fl (30).

**Figure 1.**
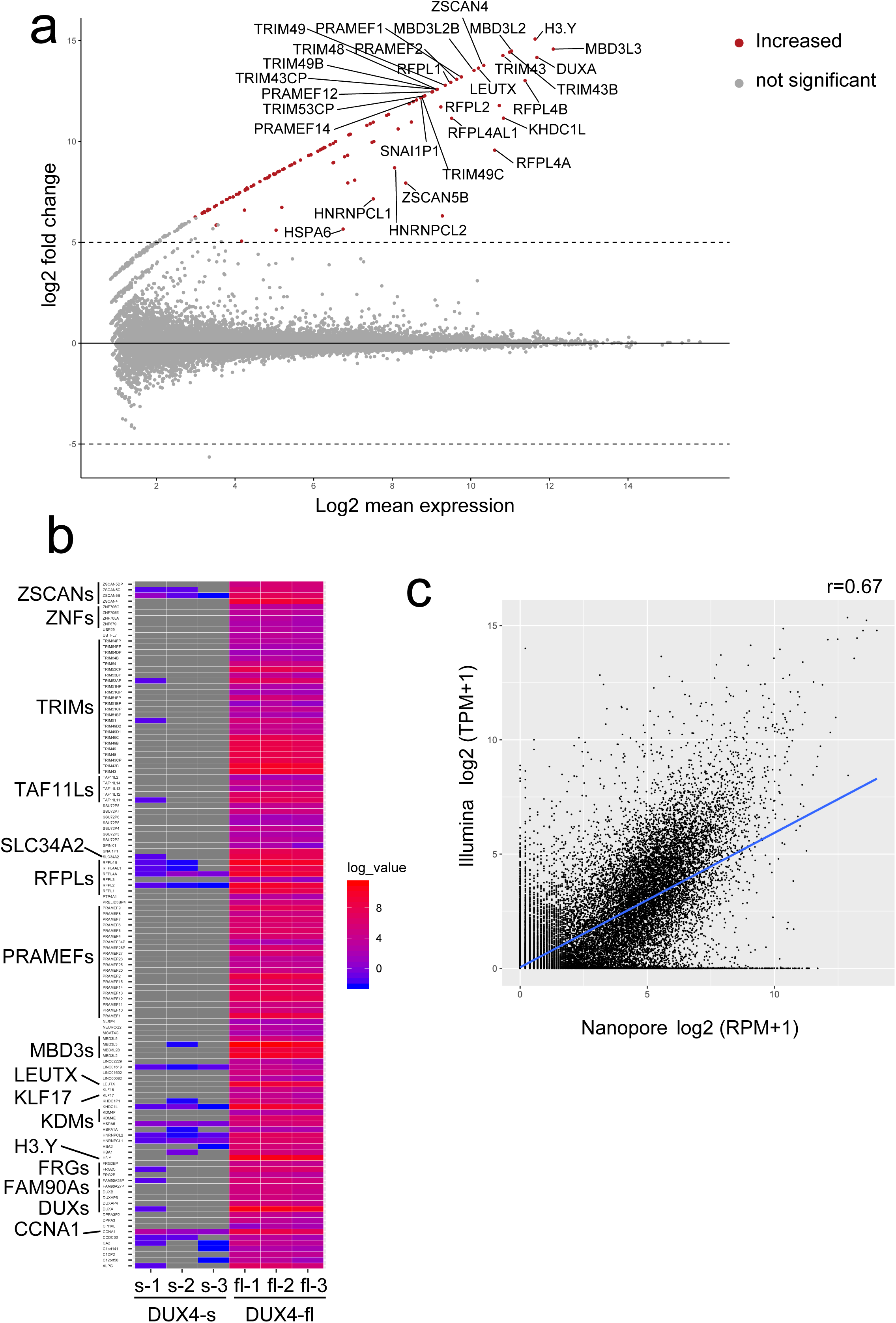
(a) MAplot for differentially expressed genes (DEG) in triplicated nanopore dRNA-seq data from DUX4-fl and DUX4-s overexpressing RD cells. Only the top 30 genes are shown. The transcripts with adjusted p-value <0.001 and log2fold change>=5 were determined as differentially expressed transcripts. (b) Heatmap of DEG genes. Well known DUX4 target genes were significantly upregulated by DUX4-fl overexpression. (c) Correlation between Illumina and Nanopore dRNA-seq expression in DUX4-fl induced genes. Pearson correlation coefficients are shown.

Next, we counted transcripts from repetitive sequence by dRNA-seq, using RepeatMasker with two different annotations: Repbase (version Open-3.0, downloaded from the UCSC genome browser, https://genome.ucsc.edu) and Dfam (version 3.1, https://www.dfam.org). We then estimated differentially expressed repeats (DERs) using DESeq2 (Figure 2a, b, Supplementary Figure 6a, b). Note that a DER represents transcripts that overlap one repeat locus. Most DERs (excluding tandem repeats) have overlapping Repbase and Dfam annotations, so we merged these annotations (Supplementary Table 4). We checked overlap between Repbase and Dfam, and found 44 loci of 61 total DERs agreed (Figure 2c). As tandem repeat annotation differs in the two databases (e.g. Repbase does not have HSATII annotation), we listed them separately (Supplementary Table 5). LTR retrotransposons were significantly enriched as previously reported, as well as satellite repeats (Supplementary Figure 7, Supplementary Tables 6, 7) (10). RT-PCR analysis confirmed four representative DERs induced by DUX4-fl (Supplementary Figure 8, Primer sequences are in Supplementary Table 8), which had no or very low expression in DUX4-s transfected cells. Publicly available Illumina short-read data for repeats did not have good correlation with nanopore dRNA-seq, probably because short reads have difficulty in detecting the whole transcripts (Figure 2d, Supplementary Figure 9).

**Figure 2.**
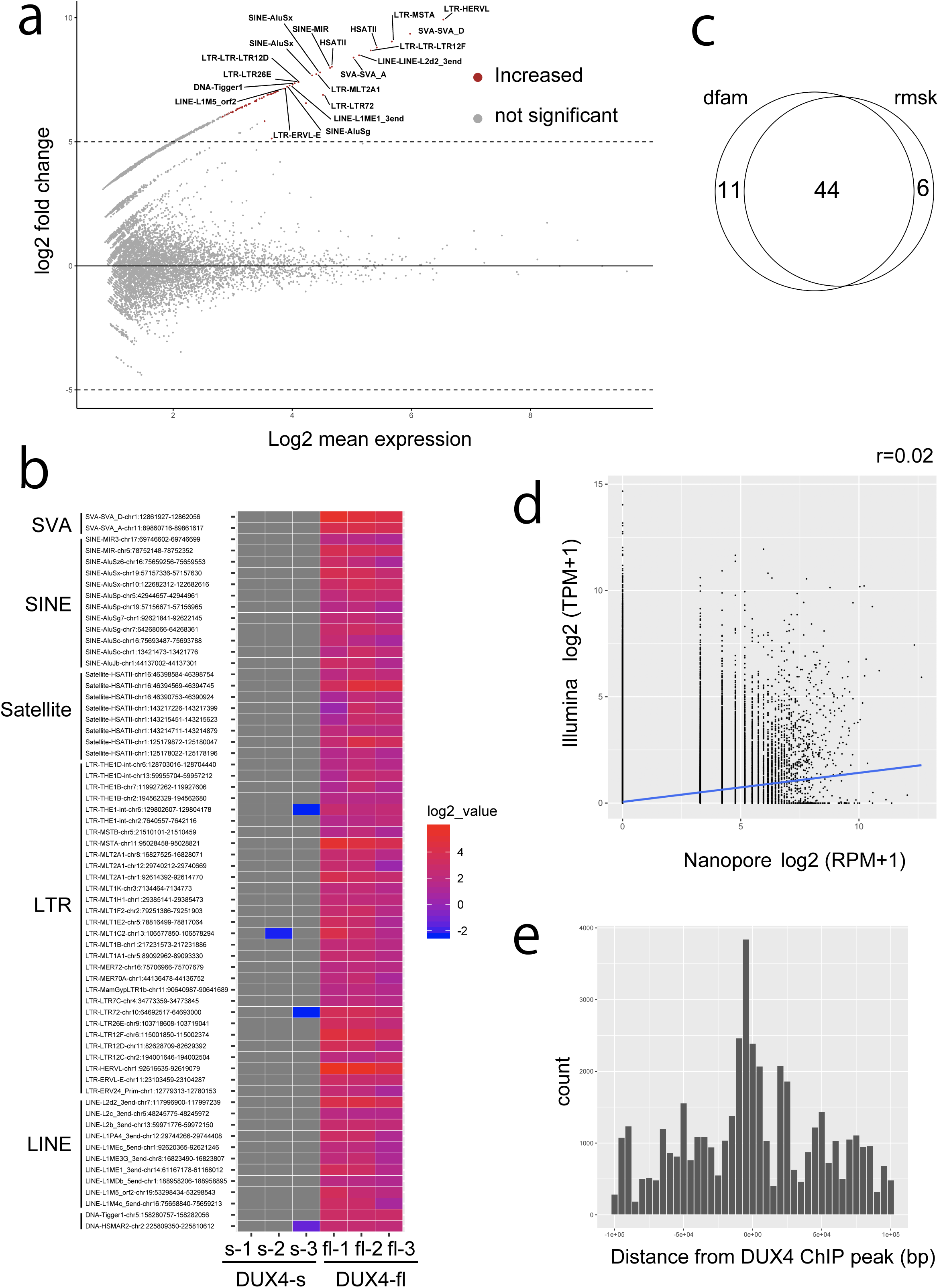
(a) MAplot for differentially expressed repeats (DER) from Dfam repeats in triplicated dRNA-seq data from DUX4-fl and DUX4-s overexpressing RD cells. Only the top 20 repeats are shown. Transcripts from repetitive elements with adjusted p-value <0.001 and log2fold change>=5 were determined as differentially expressed repeats. (b) Heatmap of DER. Many of the upregulated DERs are ERV-MaLR, which confirms previous reports. (c) Venn diagram shows 44 repeats overlap in the Dfam and Repbase annotations. (d) Correlation is not observed (r=0.02) between nanopore and Illumina read counts in Dfam repeats. Pearson correlation coefficients are shown. (e) DER repeats were located near the DUX4 ChIP peaks previously reported (10).

We also compared the detected DER loci with published DUX4-ChIP peaks (10). DUX4-ChIP peaks were near these DERs (Figure 2e, Supplementary Figure 6c), suggesting DERs are actually activated by DUX4-fl.

It was reported that DUX4-fl induces pericentric human satellite repeats HSATII bidirectionally and one strand is less abundant than the other (17). Our dRNA-seq showed that expression of satellite repeats from five loci was induced by DUX4-fl. Among them, we confirmed the expression of HSATII from chromosome 1 as previously reported (17). The satellite repeats from the five loci were expressed almost unidirectionally, but only a few opposite-strand reads were detected (Supplementary Figure 10, Supplementary Table 5). These results suggest that DUX4-fl may regulate transcription of satellite repeats on multiple chromosomes in a mostly-unidirectional manner.

### Splicing is common in DUX4-induced ERVs

ERVs are known to be processed to produce mature transcripts (31). We found most of the DUX4-induced ERVs were spliced (Figure3a, b, Supplementary Figure 11) using the canonical splice donor and acceptor signals (Figure 3c, Supplementary Figure 12), although the precise coordinates of the splice sites are uncertain because of error-prone nanopore sequencing (Supplementary Figure 2, Supplementary Table 1). Furthermore, we found that non-LTR retrotransposons (e.g. Alu) are actually transcribed from nearby LTRs and processed. Long-read sequence revealed that 35 of 61 DERs were spliced (Supplementary Table 4). We also found that 12 DERs were located downstream of DUX4-induced genes and six DERs were located within introns of the genes.

**Figure 3.**
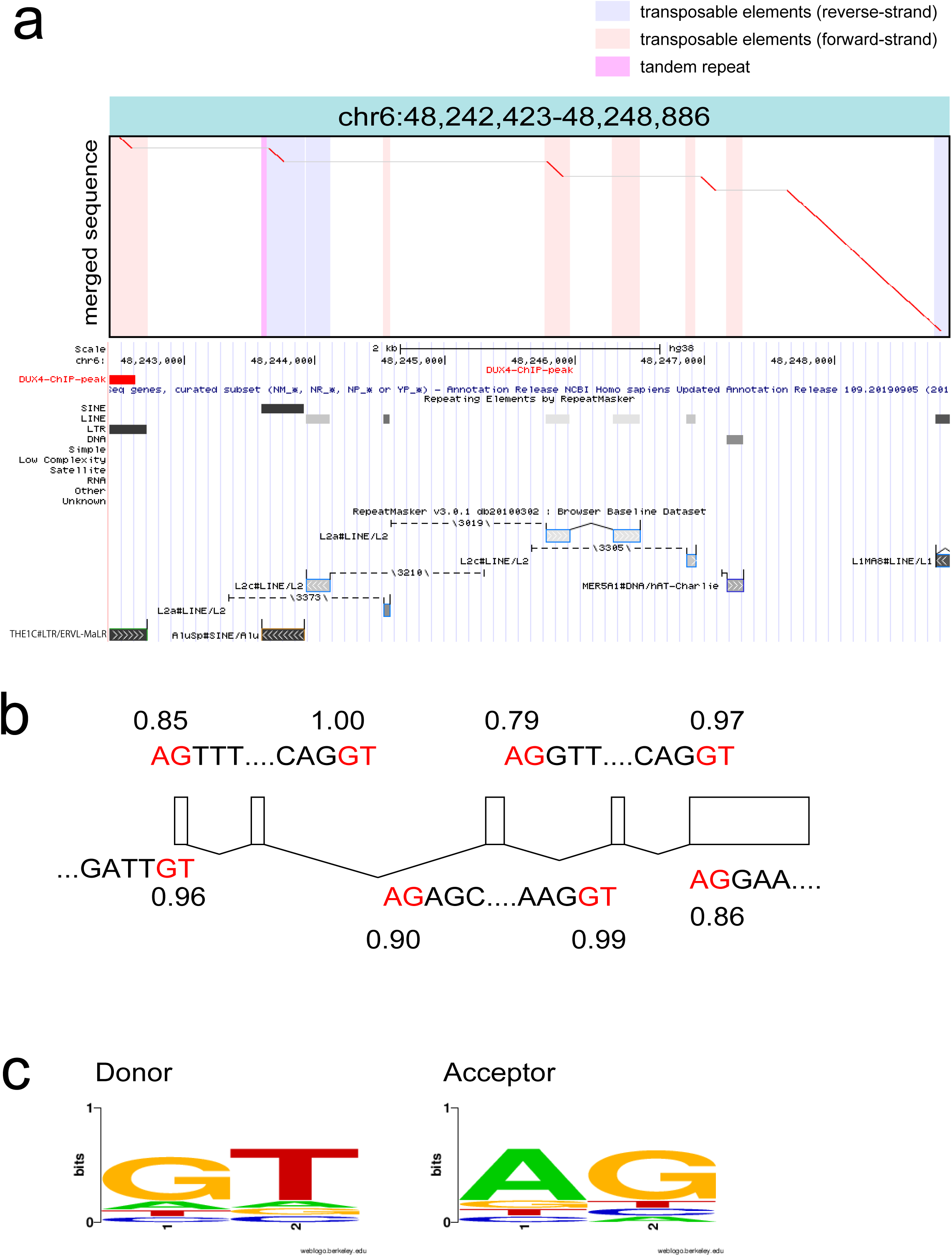
(a) An example dotplot of spliced repeat transcript. All detected transcripts from fl-1, fl-2 and fl-3 were merged using lamassemble, then aligned to the reference genome. This transcript was transcribed from an ERVL-MaLR, and then 4 splicing events occurred. Merged sequence is on the vertical axis and reference genome is on the horizontal axis in the dotplot. Vertical colored striped represent repeats in the reference genome. Below the dotplot, RepeatMasker with Repbase annotation is shown from the UCSC genome browser. (b) Splicing occurred at canonical splice donor and acceptor signals (red letters). Splice site prediction was done using NNSPLICE (v.9.0) (https://www.fruitfly.org/seq_tools/splice.html). The splice donor and acceptor prediction scores are shown. All splicing sites were predicted from LAST alignment. (c) Donor and acceptor sequences are shown as a sequence logo. Sequence logo was created by WebLogo (https://weblogo.berkeley.edu/logo.cgi).

### Fusion transcripts with LTR retrotransposons

The LTRs of DUX4-fl induced ERVs were reported to act as promoters and to generate fusion transcripts of ERVs with nearby genes. We explored fusion transcripts including LTRs (called “LTR fusion transcripts”) using dRNA-seq. As one of the triplicated DUX4-fl replicates (fl-1) has less data than the other two replicates, we only listed LTR fusion transcripts that are present in both of two replicates (fl-2 and fl-3) and never present in any of the DUX4-s overexpressing replicates. We listed 247 such LTR fusion transcripts with genes, pseudogenes or lncRNAs (Supplementary Figure 13, Supplementary Table 9), among which 216 were not reported previously (14). Some examples of DUX4-fl induced LTR fusion transcripts are shown in Figure 4a, and Supplementary Figure 12. Among the 247 LTR fusion transcripts, 15 transcripts showed statistically significant increase of gene body expression (Fig. 4b). Although we confirmed 25 LTR fusion transcripts that were reported by Young et al. (14), we could not detect 86 LTR fusion transcripts. Most of these transcripts only have <100 raw Illumina reads, which may be hard to detect with smaller numbers of nanopore dRNA-seq reads (Supplementary Figure 14). Interestingly, the long read sequencing detected splicing variants of some of the LTR fusion transcripts. For example, the previous study reported RBM26 as one LTR fusion transcript (Supplemental Figure 9g). However, our analysis revealed that two different splicing isoforms of the LTR-RBM26 fusion transcript are induced by DUX4-fl. This result suggests that long read sequencing has an advantage for detecting whole isoforms of transcripts.

**Figure 4.**
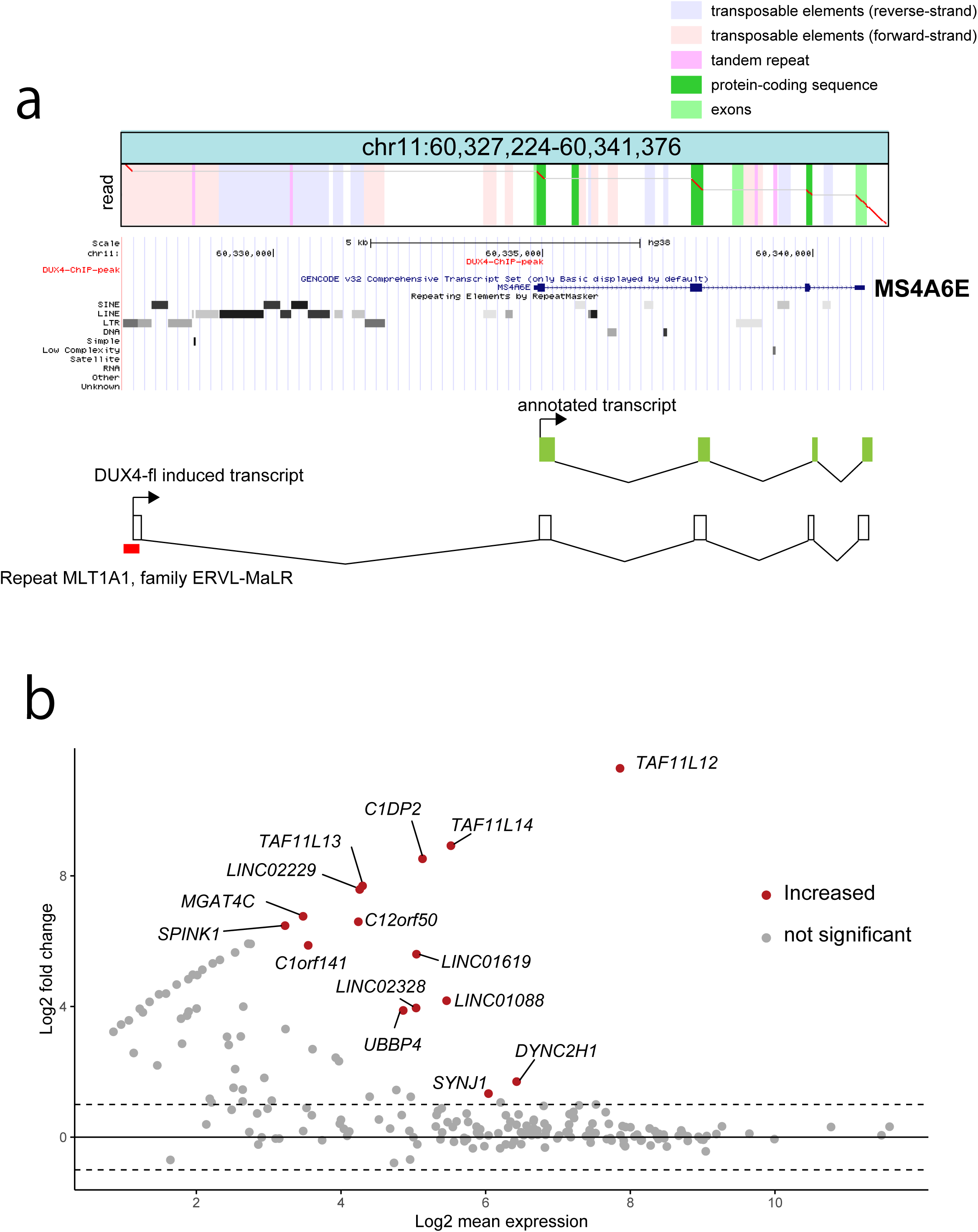
(a) An example fusion transcript read of an ERVL-MaLR with the MS4A6E gene, induced by DUX4-fl. Below the dotplot, RepeatMasker with Repbase annotation is shown from the UCSC genome browser. (b) MAplot of DEG of fusion transcripts from DUX4-activated repeats. Fifteen fusion transcripts (genes) show significant upregulation compared to DUX4-s transfected cells.

## Discussion

Direct sequencing of RNA molecules is a highly anticipated technology because it avoids PCR or reverse transcription biases and also has the possibility to provide RNA modification signatures. However, it is still not clear whether nanopore dRNA-seq is suited for much biology research because of the requirement for a large amount of RNA and low throughput (23). Our study showed that dRNA-seq using PromethION, a high-throughput nanopore sequencer, is applicable to detect differentially expressed transcripts including genes, lncRNA, satellite repeats, and retrotransposons, and also provides example datasets to the research community.

The transcriptome of DUX4-fl expressing myoblasts using an Illumina short read sequencer was previously reported (10, 14). We compared our results using nanopore dRNA-seq to the previous findings. As we did not do Illumina sequencing ourselves and utilized public data which has no biological replicates, fair comparison may be difficult. Nevertheless, we saw some consistency between genes expressed in nanopore and Illumina data. We observed activation of known DUX4-induced germline genes and cell cycle genes in our dRNA-seq data, comparable to the previous reports. In addition, activation of ERVs and tandem repeats are also comparable to the previous results, which shows the feasibility of using dRNA-seq to detect DUX4-activated repeats. We identified 79 significantly DUX4-induced repetitive elements (including 18 tandem repeats, which may be overlapping), using two different repetitive sequence databases (Repbase and Dfam). Most of them were transcribed from DUX4-activated LTRs, as reported previously (14). In addition to the known findings, long read dRNA-seq clearly shows that RNA splicing is a common event in DUX4-activated ERV transcripts.

ERVs have two long-terminal repeat (LTR) sequences that actually have promoter activity, and some of them are co-opted as promoters affecting the expression of nearby genes (32). It has been reported that binding of DUX4 to the LTRs activate genes or lncRNA. Our dRNA-seq identified novel transcripts in addition to reported ones. DESeq2 analysis confirmed that some of these fusion genes are actually upregulated compared to DUX4-s, suggesting that these DUX4-fl-activated aberrant transcripts contribute to altered gene expression (Figure 4b). We also identified several pseudogenes such as *UBBP4* that were activated by DUX4. These pseudogenes may actually be genes, and can only be detected in rare situations (e.g. cleavage-stage embryo or FSHD muscle).

It was reported that DUX4-fl also activates satellite repeats and causes toxic intranuclear dsRNA foci (17). Our dRNA-seq also detected several satellite repeat loci. We did not observe obvious bidirectional transcription of satellite repeats: only a few antisense strand reads were observed, which may be concordant with a previous report (17). Nanopore sequencing error in reading the opposite strand seems an unlikely explanation, because several different tandem repeats showed a unidirectional pattern.

Our results show that most of the DUX4-activated non-LTR retrotransposons (e.g. Alu) were transcribed from ERVs and underwent RNA processing, indicating the importance of analyzing full-length transcripts. Although Alu is known to form dsRNAs, our dRNA-seq analysis did not detect inverted tandem Alu that is often contained in human dsRNA.

In conclusion, by nanopore dRNA-seq with a PromethION, we obtained enough reads to detect DUX4-fl-induced genes and transposons with statistical significance. In addition, we could detect novel transcripts that were activated by DUX4-bound LTRs. Further improvements of the technologies (e.g. reducing the required input RNA amount or increased output) may expand the usefulness of this technology.

## Methods

### Cell culture, DUX4 transfection and extracting RNA

Human rhabdomyosarcoma-derived cell line, RD cells were obtained from the ATCC (CCL-136, University Boulevard Manassas, VA, USA). The RD cells were cultured in DMEM (cat. D5796, Sigma-Aldrich, St Louis, MO) supplemented with fetal bovine serum (FBS; cat. 10270-106, Thermo-Fisher, Grand Island NY) at 37°C in a humidified atmosphere of 5% CO_2_. The DUX4-fl and DUX4-s plasmids were prepared as described previously (5). The RD cells were plated on 6-well plates at the density of approximately 30 ⨯ 10^4^ cells per well. The next day, 2 g of DUX4 expression constructs were transfected into RD cells with 6 L of the X-treme GENE 9 DNA transfection reagent (cat. XTG9-RO, Roche Diagnostics, Indianapolis, USA) diluted in 100 L of Opti-MEM I (Gibco) following the manufacturer’s instructions.

24h after transfection, total RNA was extracted with TRIzol reagent (cat. 15596018, invitrogen) with DNase I treatment (cat.79254, QIAGEN). The total RNA solution from 12 wells are passed through one column of RNeasy mini kit (cat.74104, QIAGEN) to obtain approximately 100 g of purified total RNA. Then, poly-A RNA was isolated from 60∼90ug total RNA using μMACS mRNA Isolation Kits (Miltenyi Biotec Inc. CA, USA). Poly-A RNA (500 ng) was subjected to nanopore library preparation.

### PromethION direct RNA nanopore sequencing

Library preparation was done using RNA002 Direct RNA sequencing kit according to the manufacturer’s protocol and sequenced by PromethION sequencer using one flowcell per one sample (Oxford Nanopore Technologies, UK). Basecalling was done by MinKNOW with default settings.

### Mapping to the human reference genome

Nanopore direct RNA sequence reads (dRNA-seq.fastq) were mapped to the human reference genome GRCh38 according to the online instructions (https://github.com/mcfrith/last-rna/blob/master/last-long-reads.md).

Briefly, a reference database was generated as follows:

lastdb -P8 -uNEAR -R01 GRCh38 GRCh38.fa

Then, nanopore reads were aligned to GRCh38 with parameters for each dataset

obtained using last-train.

last-train -Q0 GRCh38 dRNA-seq.fastq > train.out

parallel-fasta “lastal -p train.out -d90 -m20 -D10 mydb | last-split -d2 -g GRCh38” < dRNA-seq.fastq > aln.maf

Note the -d2 option is for cDNA of unknown/mixed strands. dRNA-seq reads are presumably of RNA forward strands; this option may be omitted.

### Finding and counting gene and repeat overlapping reads

Aligned files (aln.maf) were subjected to read counting according to the online instructions (https://github.com/mcfrith/last-rna/blob/master/scripts.md).

Gene annotation (knownGene.txt) was obtained from the UCSC genome browser (http://hgdownload.soe.ucsc.edu/goldenPath/hg38/database/).

First, gene-overlap regions were obtained like this:

gene-overlap.sh genes file.maf > out

tail -n+2 out | cut -f3 | cut -d, -f1 | uniq -c | awk ‘{print $2” \t” $1}’ | sort > gene-count For counting repeat regions, the above gene annotation was combined with Repbase (rmsk.txt) or Dfam annotation, and then only repeat regions were examined.

gene-overlap.sh genes+repeats file.maf > out

tail -n+2 out | cut -f3 | uniq -c | grep ‘rmsk,’ | awk ‘{print $2” \t” $1}’ | sort > repeat-count

### Merging nanopore reads using lamassemble

Nanopore reads were merged into consensus sequence using lamassemble (33, 34). We used fl-2 last-train output as a representative parameter set.

lamassemble train-out merged.fa > consensus.fa

The consensus sequence was realigned to GRCh38 and then dotplot pictures were drawn as previously described (33).

### Differentially expressed genes or repeats

We estimated differentially expressed genes or repeats using DESeq2 (28). For both genes and repeats, we determined the transcripts with adjusted p-value <0.001 and log2fold change>=5 as differentially expressed transcripts.

### Repeat enrichment analysis

We conducted an enrichment analysis of each repeat name (Dfam and Repbase), class and family (Repbase) of DERs by comparing the observed fraction of repeats with the fraction identified in the human genome (GRCh38). Based on those fractions and the number of repeats, Z-scores were calculated, and adjusted p-values were also obtained based on chi-squared tests with Bonferroni correction.

### Finding fusion transcripts with LTRs

To find fusion transcripts of genes with LTRs, we used the gene-fusion script from: (https://github.com/mcfrith/last-rna/blob/master/scripts.md).

gene-fusions.sh genes.txt alignments.maf > fusions.txt

Transcripts fused with LTR annotations from RepeatMasker+Repbase that are upstream or in the gene bodies are counted.

As the number of reads in one of the DUX4-fl replicates was about half of the other two replicates, we counted only the reads that were not expressed any of three replicates with DUX-s expression. Differentially expressed fusion transcripts were also determined by DESeq2.

### Public Illumina cDNA seq and ChIP seq data

Publicly available Illumina short read cDNA seq data of DUX4 overexpressing human myoblast cells (MB-135, DUX4-fl or GFP was introduced by lentiviral infection.

Accession numbers are SRR823337 and SRR8233378) were re-analyzed using STAR v2.7 (35). Ribosomal RNA sequence was removed using bowtie2 v2.2.4 (36) with iGenome indexes (https://jp.support.illumina.com/sequencing/sequencing_software/igenome.html). Only uniquely mapped reads (MAPQ 255) were counted.

Coordinates of DUX4 ChIP-seq peaks (10) were converted to GRCh38 using UCSC liftOver (https://genome.ucsc.edu/cgi-bin/hgLiftOver). We used the middle coordinate of the peak and that of the repeat to show the distance.

## Acknowledgements

This work was supported by JSPS KAKENHI under the grant numbers JP19K07977 and 16H06279 (PAGS) (to S. Mitsuhashi), 16H06429, 16K21723, 17H05823, and 20K06775 (to S. N.), and 18K07511 (to H. Mitsuhashi). Computations were partially performed on the NIG supercomputer at ROIS National Institute of Genetics and partially performed on Matsumoto-lab computer.

## Author contributions

S.M. and H.M contributed to acquisition and analysis of the data. All authors contributed to the conception of the work and interpretation of the data, and read and approved the final manuscript.

## Competing interests

The authors declare no competing interests.

## Data availability

Direct RNAseq data is available under accession numbers DRA008398 (under project number PRJDB8318) from DDBJ DRA database (https://www.ddbj.nig.ac.jp/dra/index-e.html). Illumina human myoblast sequences were obtained under accession numbers SRR823337 and SRR823338.

## Web resources

UCSC genome browser:

LAST: http://last.cbrc.jp

Long read alignment using LAST:

https://github.com/mcfrith/last-rna/blob/master/last-long-reads.md

lamassemble: https://gitlab.com/mcfrith/lamassemble

STAR: https://github.com/alexdobin/STAR

Illumina iGenome:

https://support.illumina.com/sequencing/sequencing_software/igenome.html

## Notes

### Competing Interest Statement

The authors have declared no competing interest.

