## Supplementary Figures for "Nanopore direct RNA sequencing detects DUX4-activated repeats and isoforms in human muscle cells"

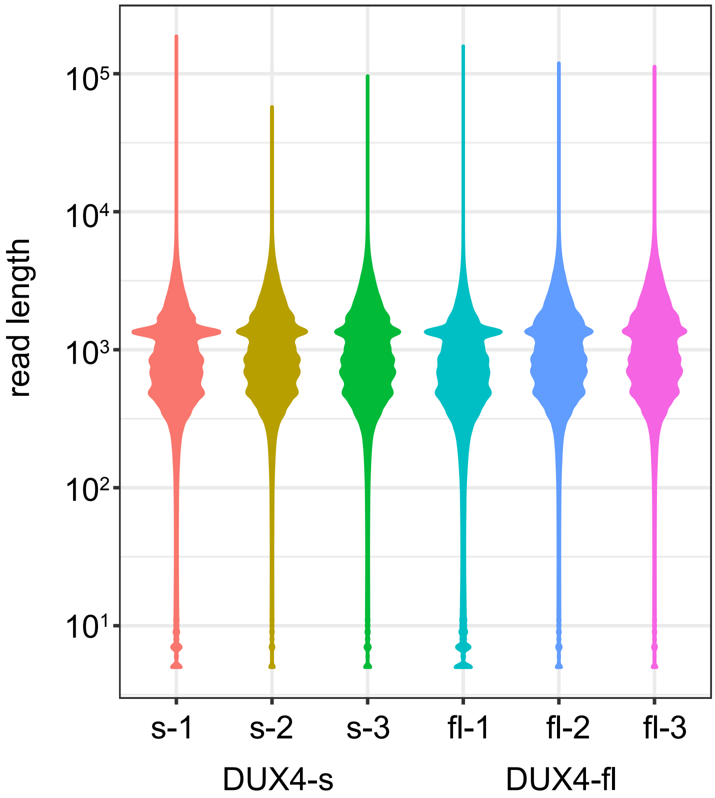


Supplementary Figure 1.
Violin plot for the read length of 6 samples (DUX4-fl; fl-1, fl-2 and fl-3. DUX4-s; s-1, s-2 and s-3. All showed similar distribution patterns.

| | A | C | G | T |
|---|---|---|---|---|
| A | 0.25 | 0.0015 | 0.0070 | 0.0077 |
| C | 0.0018 | 0.22 | 0.00035 | 0.015 |
| G | 0.0067 | 0.00039 | 0.24 | 0.00068 |
| T | 0.0079 | 0.0128 | 0.00062 | 0.23 |

| | Deletion | Insertion |
|---|---|---|
| Opening probability | 0.058 | 0.030 |
| Extension probability | 0.44 | 0.38 |

Supplementary Figure 2
Rates (probabilities) of substitutions, deletions, and insertions between nanopore dRNA-seq and human reference genome GRCh38. The 4x4 matrix shows substitution probabilities. Rows correspond to genome bases and columns correspond to read bases.

a

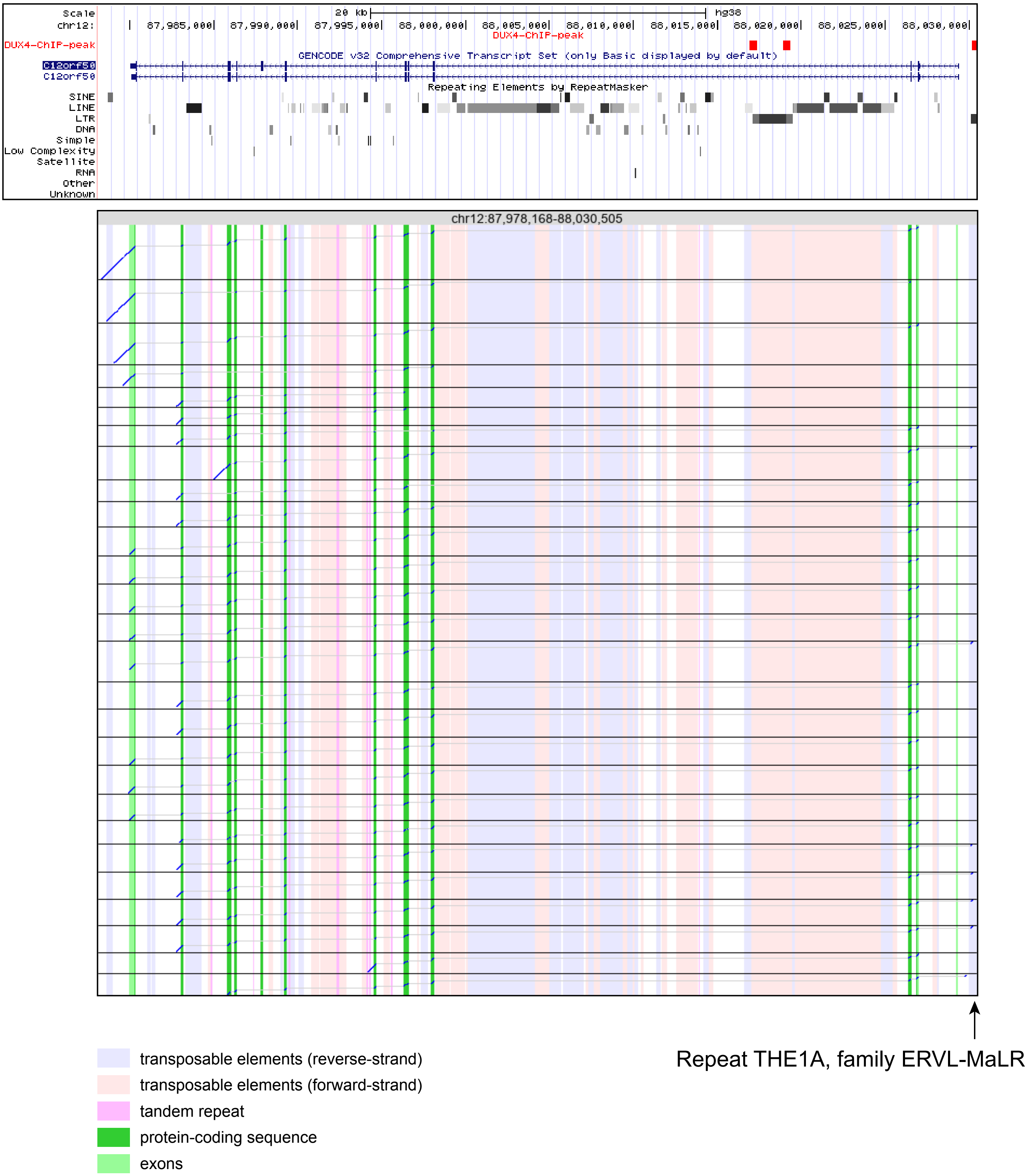
Scale
chr12:
20 kb
hg38
87,985,000
87,990,000
87,995,000
88,000,000
88,005,000
88,010,000
88,015,000
88,020,000
88,025,000
88,030,000
DUX4-ChIP-peak
GENCODE v32 Comprehensive Transcript Set (only Basic displayed by default)
C12orf50
C12orf50
Repeating Elements by RepeatMasker
SINE
LINE
LTR
DNA
Simple
Low Complexity
Satellite
RNA
Other
Unknown
chr12:87,978,168-88,030,505
Repeat THE1A, family ERVL-MaLR
transposable elements (reverse-strand)
transposable elements (forward-strand)
tandem repeat
protein-coding sequence
exons

b

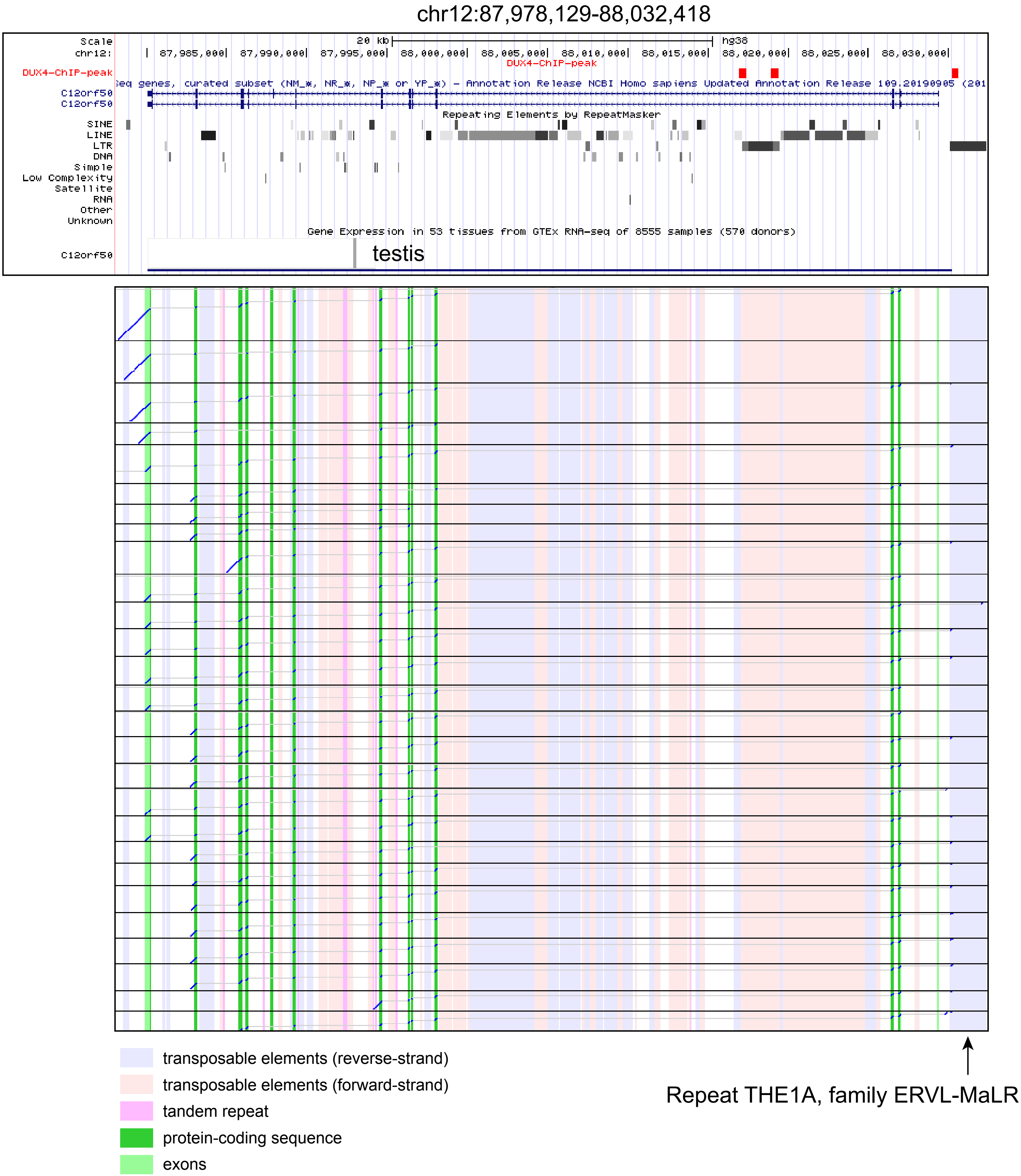


Supplementary Figure 3
Long read sequencing shows clear exon connectivity in C12orf50, which is induced by DUX4-fl. The vertical blue stripes indicate transposable elements in the reverse strand. (a) LAST alignment with default parameters. (b) LAST alignment with sensitive parameters (-d60 -m1000 -D10) shows many transcripts originate from ERVL-MaLR. Upper: UCSC genome browser view of the locus corresponding to the dotplot. C12orf50 is exclusively expressed in testis.

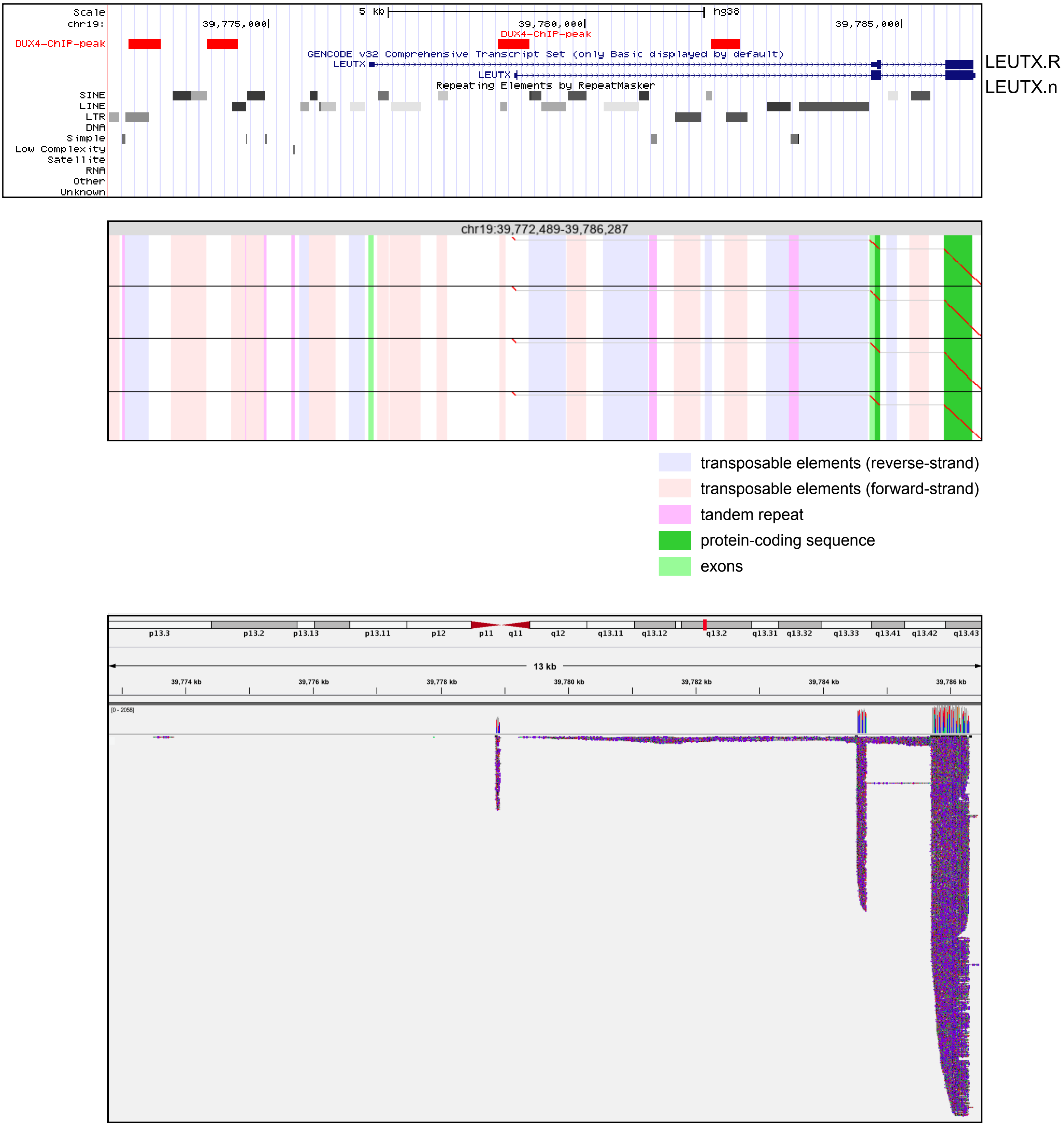


Supplementary Figure 4
Long reads show that only the functional isoform of LEUTX, LEUTX.n, is induced by DUX4-fl. The non-functional isoform LEUTX.R is not induced. Top: UCSC genome browser shows that DUX4 ChIP peak overlaps the LEUTX gene. Middle: dotplot shows 4 representative reads. Bottom: IGV picture shows read coverage from s-3 and fl-3 data.

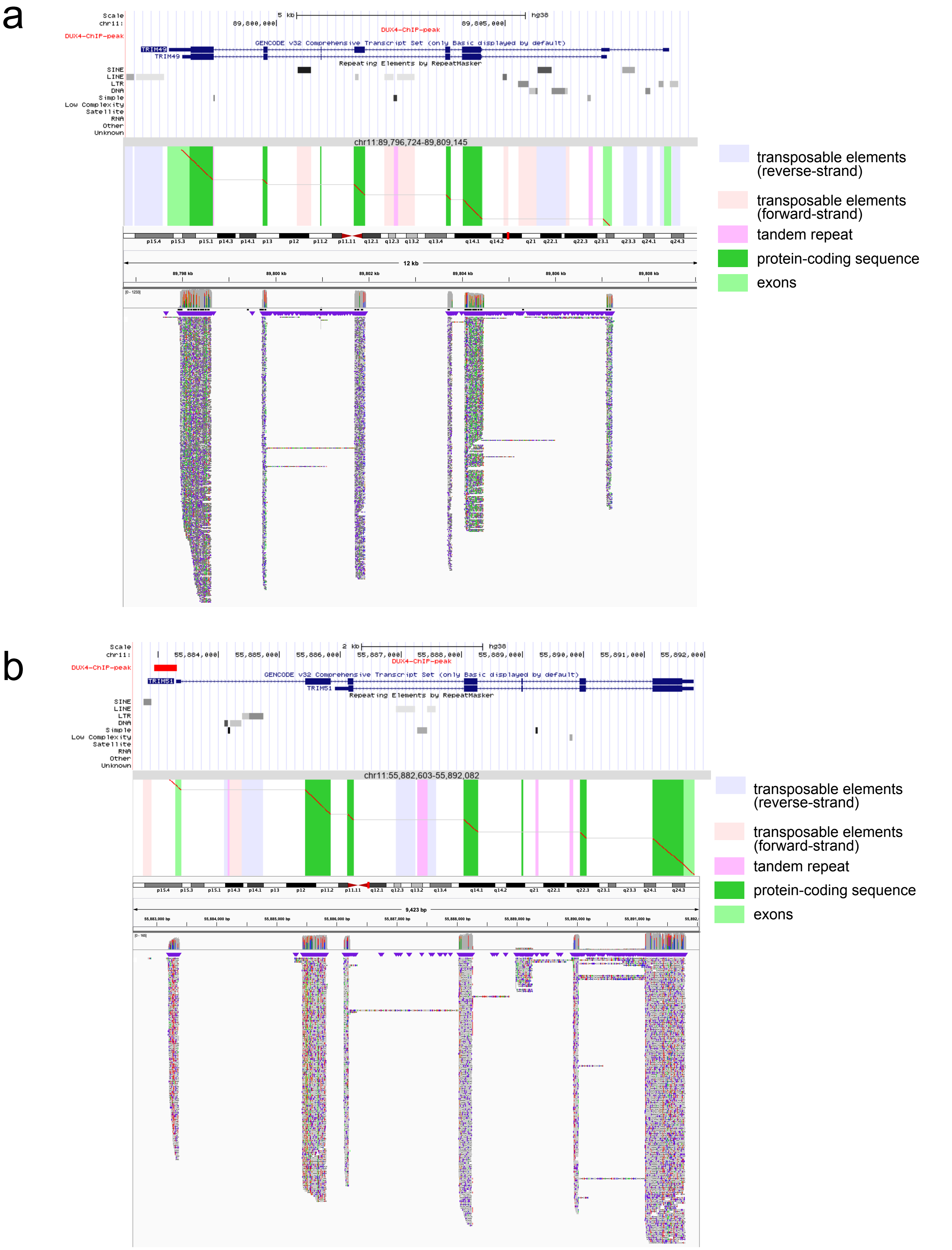


Supplementary Figure 5

Long reads show isoform-specific expression of TRIM49 (A) and TRIM51 (B). Top: UCSC genome browser shows DUX4 ChIP peak and genes. Middle: dotplot pictures of representative read. Bottom: IGV picture shows read coverage from s-3 and fl-3 data.

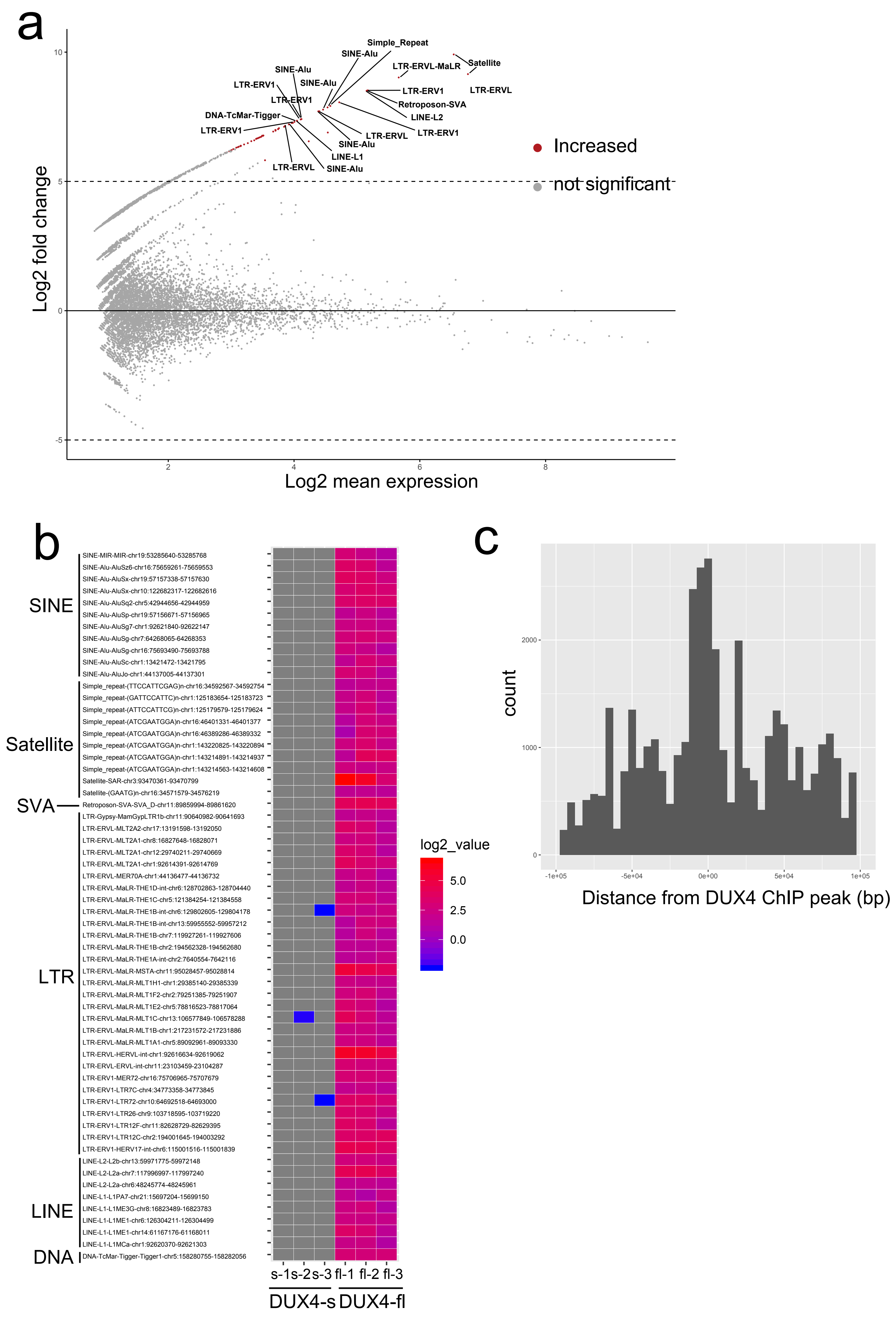


Supplementary Figure 6
(a) MAplot for differentially expressed repeats (DER) from Repbase repeats in triplicated dRNA-seq data from DUX4-fl or DUX4-s overexpressing RD cells. Only the top 20 repeats are shown.
(b) Heatmap of DER.
(c) DER were located near the DUX4 ChIP peak that was publicly available.

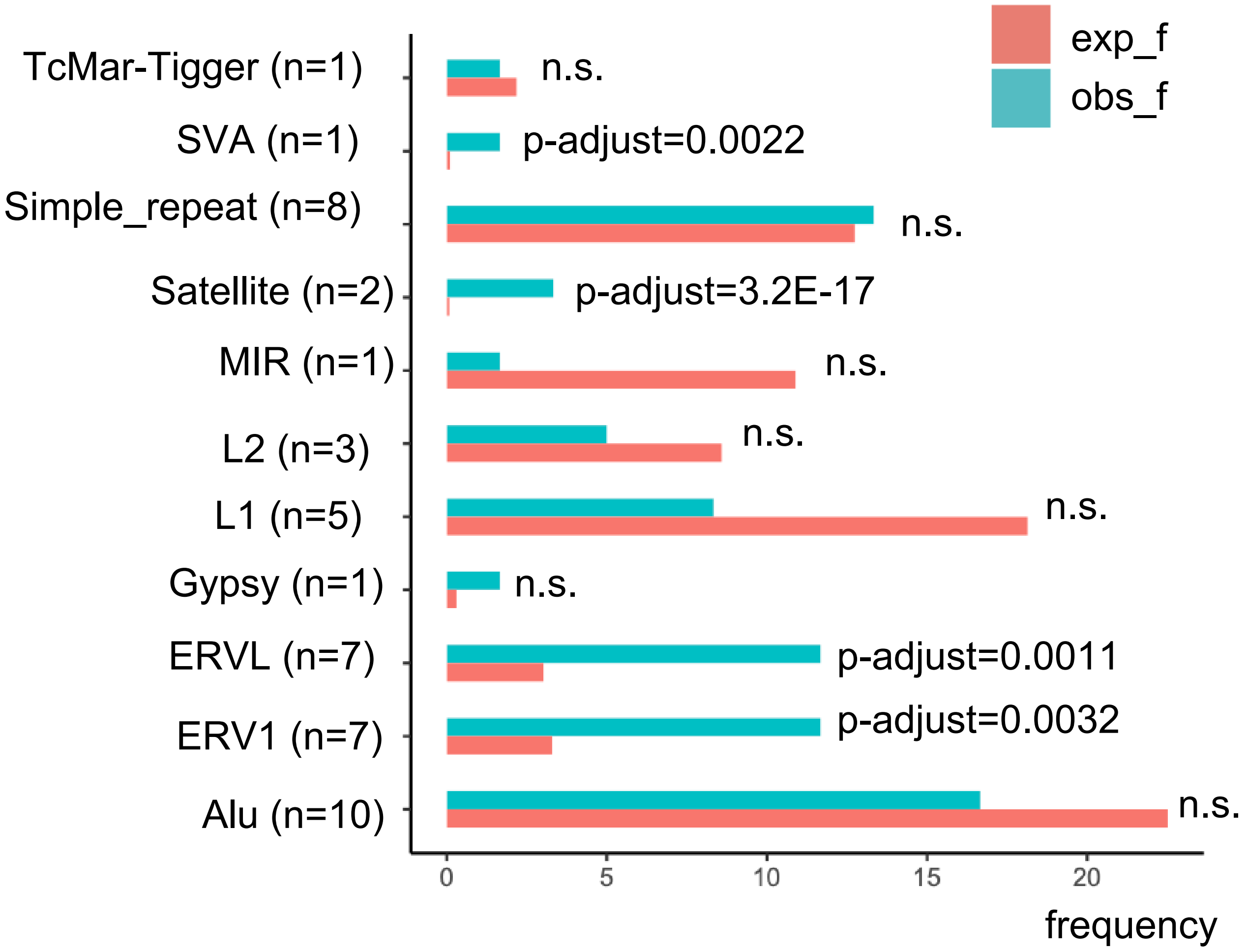


Supplementary Figure 7
ERVL-MaLR, ERV1, ERVL, Satellite and ERVK are enriched in DUX4-fl expressed DERs.
exp_f: expected frequency. obs_f: observed frequency.

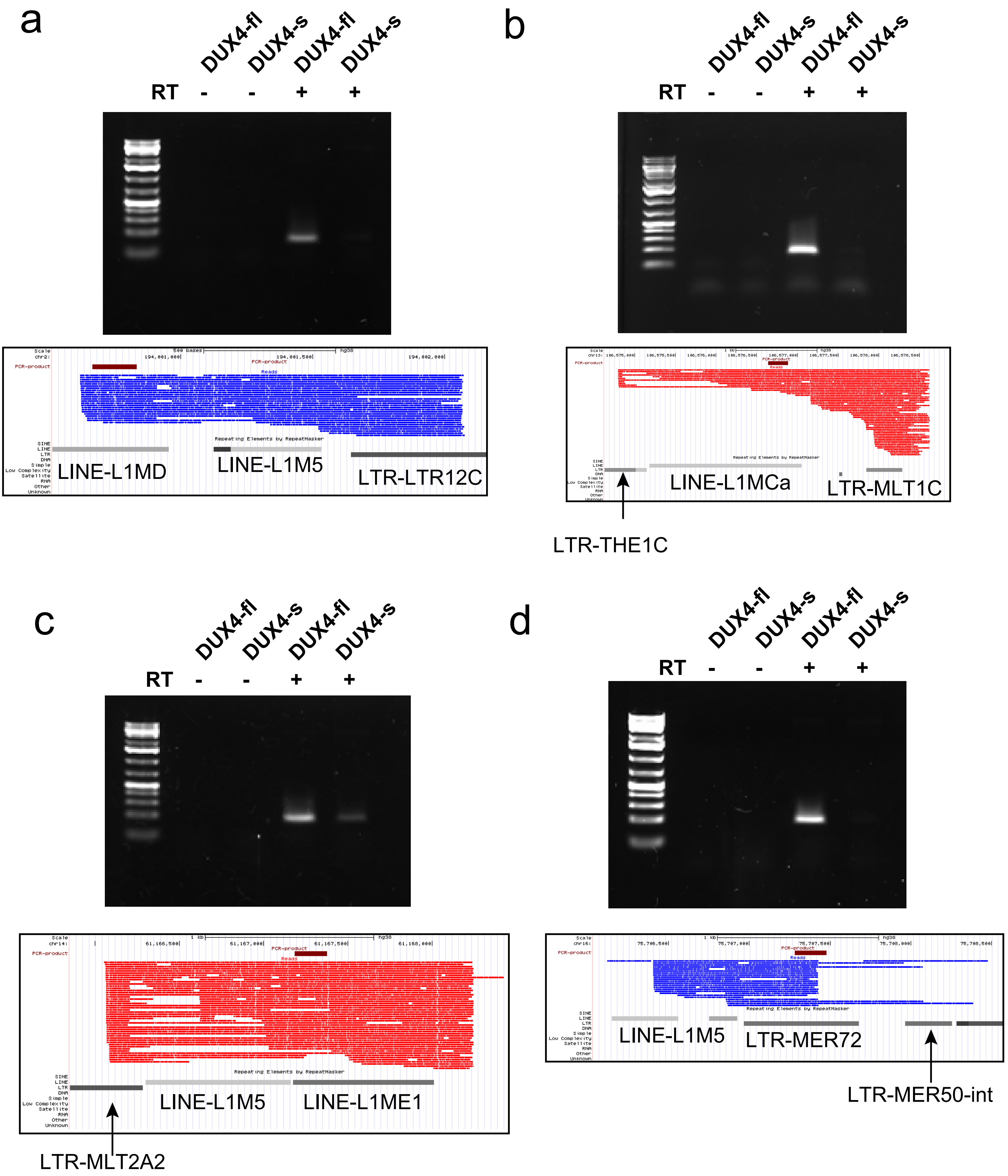


Supplementary Figure 8
Agarose gel electrophoresis of PCR products from 4 representative repeats (a, b, c, d). Primer sequence is available in Supplementary Table S8. For each repeat, nanopore reads (black bars) and PCR product location (blue bar) are shown. LAST alignment was converted to psl format using maf-convert (http://last.cbrc.jp/doc/maf-convert.html) and then visualized in UCSC genome browser. Repbase classes and names are also shown. Blue reads indicate minus direction of the transcript. Red reads indicate plus direction of the transcript. Brown bar represents the location of PCR amplicons.

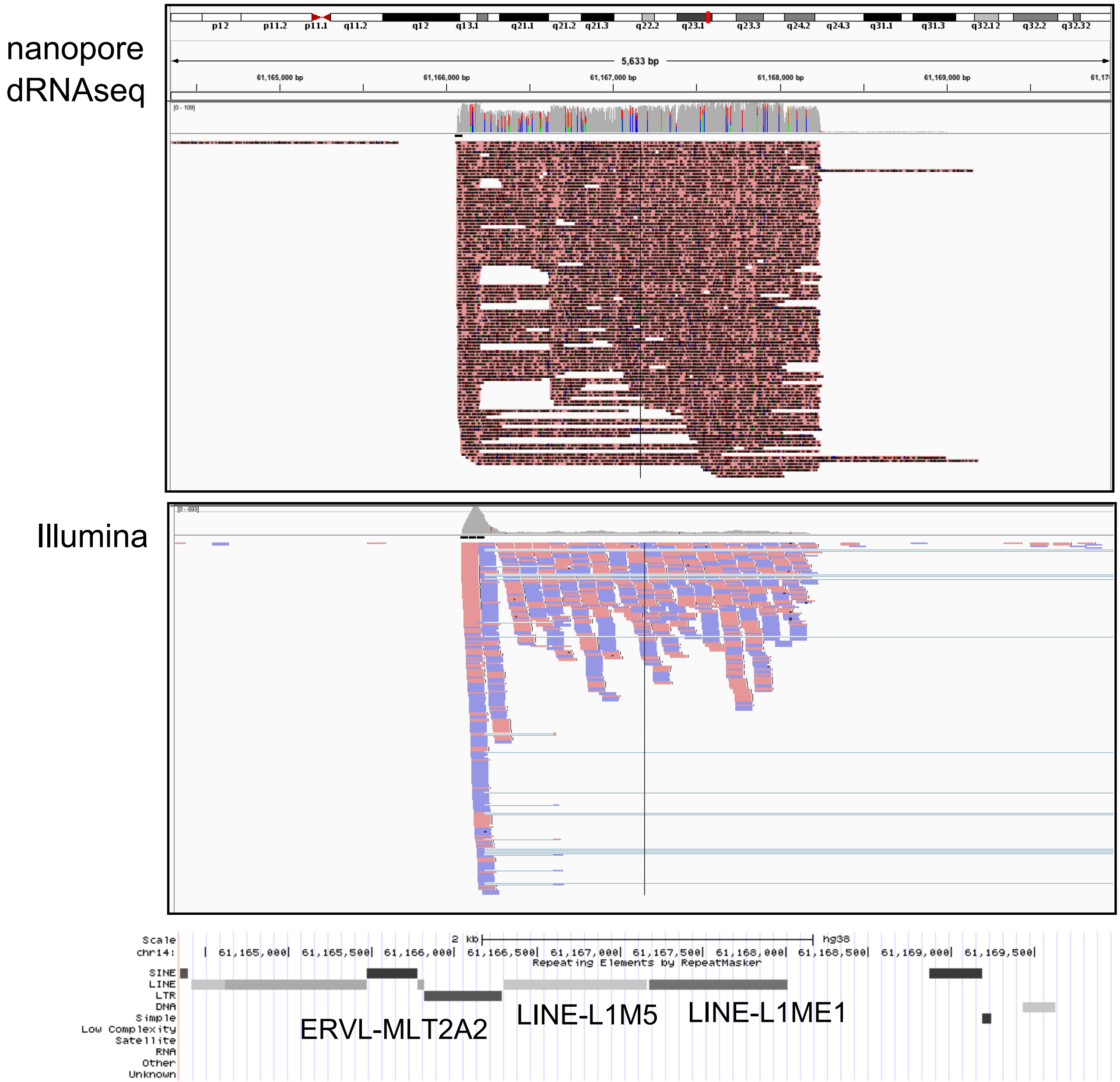


Supplementary Figure 9
IGV view of our dRNA-seq alignment and publicly available Illumina cDNA-seq (14) . LINE is transcribed from ERVL-MLT2A2. dRNA-seq reads show nearly even coverage. Illumina short reads have high coverage at ERVL-MLT2A2 but fewer reads were mapped on LINE. Only uniquely mapped reads are shown. RepeatMasker annotation from the UCSC genome browser is shown below the IGV view.

a

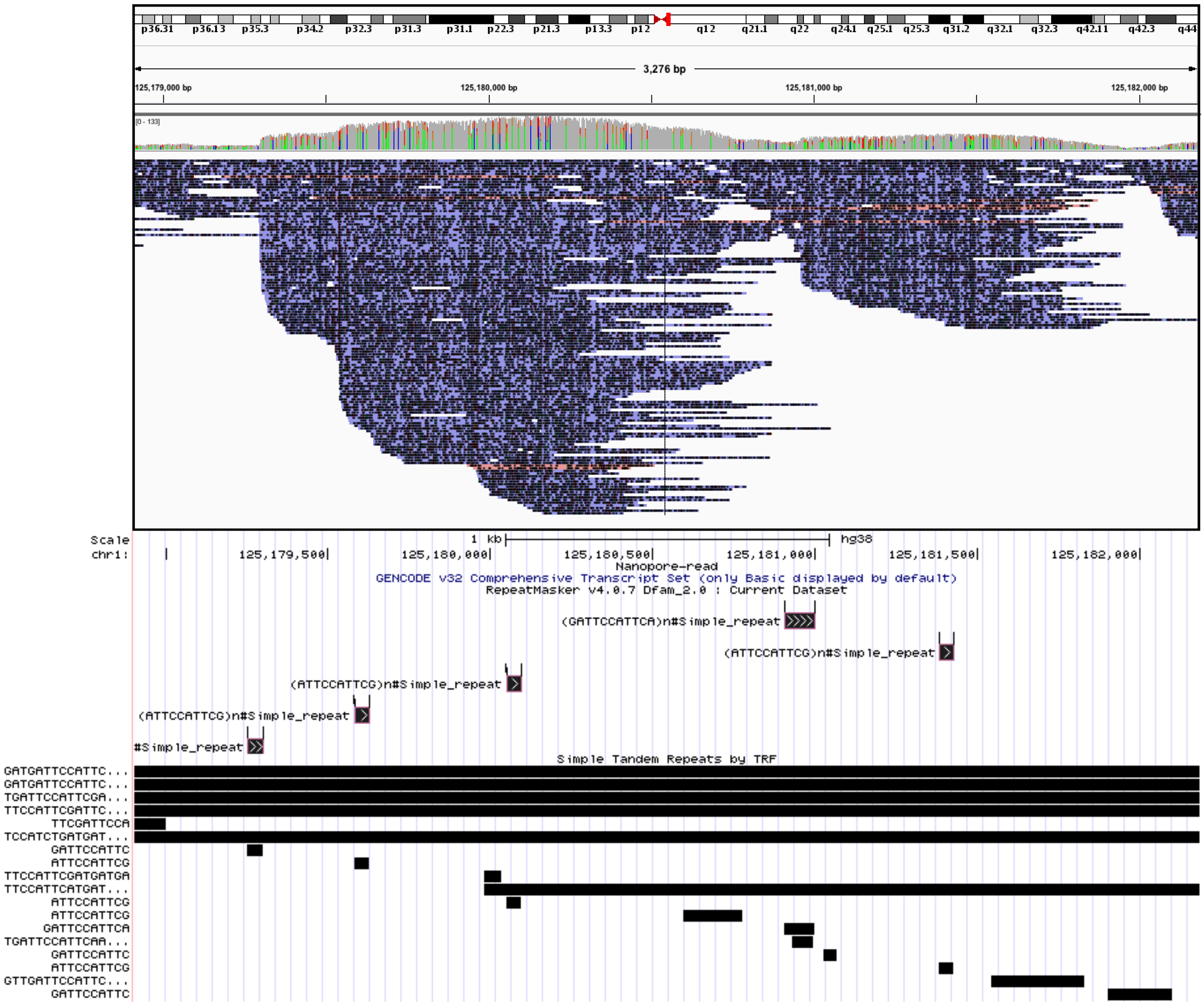
p36.31 p36.13 p35.3 p34.2 p32.3 p31.3 p31.1 p22.3 p21.3 p13.3 p12 q12 q21.1 q22 q24.1 q25.1 q25.3 q31.2 q32.1 q32.3 q42.11 q42.3 q44
3,276 bp
125,179,000 bp
125,180,000 bp
125,181,000 bp
125,182,000 bp
[0 - 133]
Scale
chr1:
1 kb
hg38
125,179,500
125,180,000
125,180,500
125,181,000
125,181,500
125,182,000
Nanopore-read
GENCODE v32 Comprehensive Transcript Set (only Basic displayed by default)
RepeatMasker v4.0.7 Dfam_2.0 : Current Dataset
(GATTCCATTCA)n#Simple_repeat
(ATTCCATTCG)n#Simple_repeat
(ATTCCATTCG)n#Simple_repeat
(ATTCCATTCG)n#Simple_repeat
#Simple_repeat
Simple Tandem Repeats by TRF
GATGATTCCATTC...
GATGATTCCATTC...
TGATTCCATTCGA...
TTCCATTCGATTC...
TTCGATTCCA
TCCATCTGATGAT...
GATTCCATTC
ATTCCATTCG
TTCCATTCGATGATGA
TTCCATTCATGAT...
ATTCCATTCG
ATTCCATTCG
GATTCCATTCA
TGATTCCATTCAA...
GATTCCATTC
ATTCCATTCG
GTTGATTCCATTC...
GATTCCATTC

b

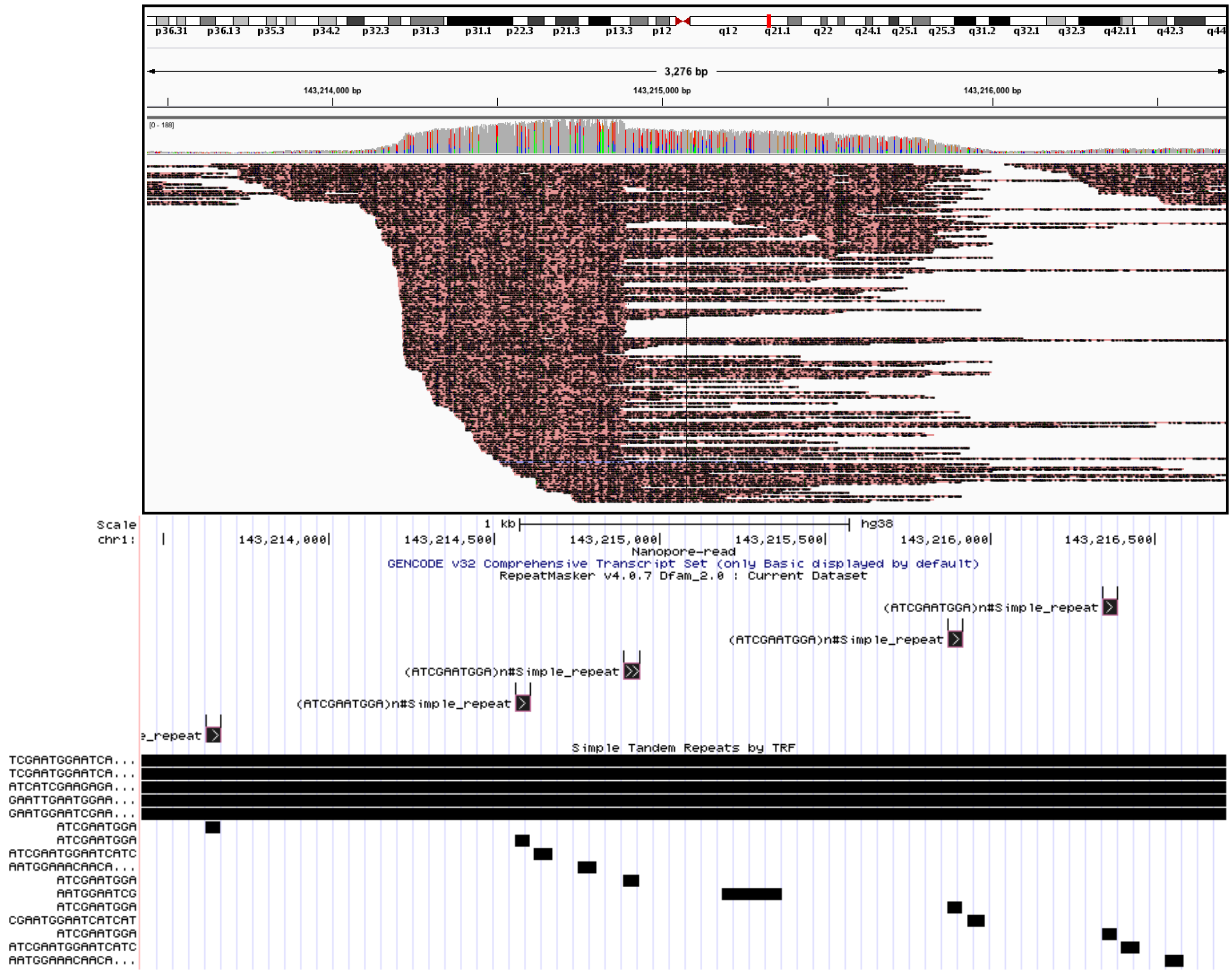


Supplementary Figure 10

(a) IGV view of a DUX4-fl-activated satellite repeat in dRNA-seq. Majority of transcripts from centromeric tandem repeat expression at chr1 (chr1:143,213,435-143,216,714) are minus stranded (shown as blue reads). A few reads are plus stranded (shown as red reads). RepeatMasker annotation from the UCSC genome browser is shown below the IGV view.

(b) IGV view of another DUX4-fl-activated satellite repeat in dRNA-seq. Majority of transcripts from another tandem repeat (chr1:125,178,904-125,182,183) are plus stranded (shown as red reads). A few reads are minus stranded (shown as red reads). RepeatMasker annotation from the UCSC genome browser is shown below the IGV view.

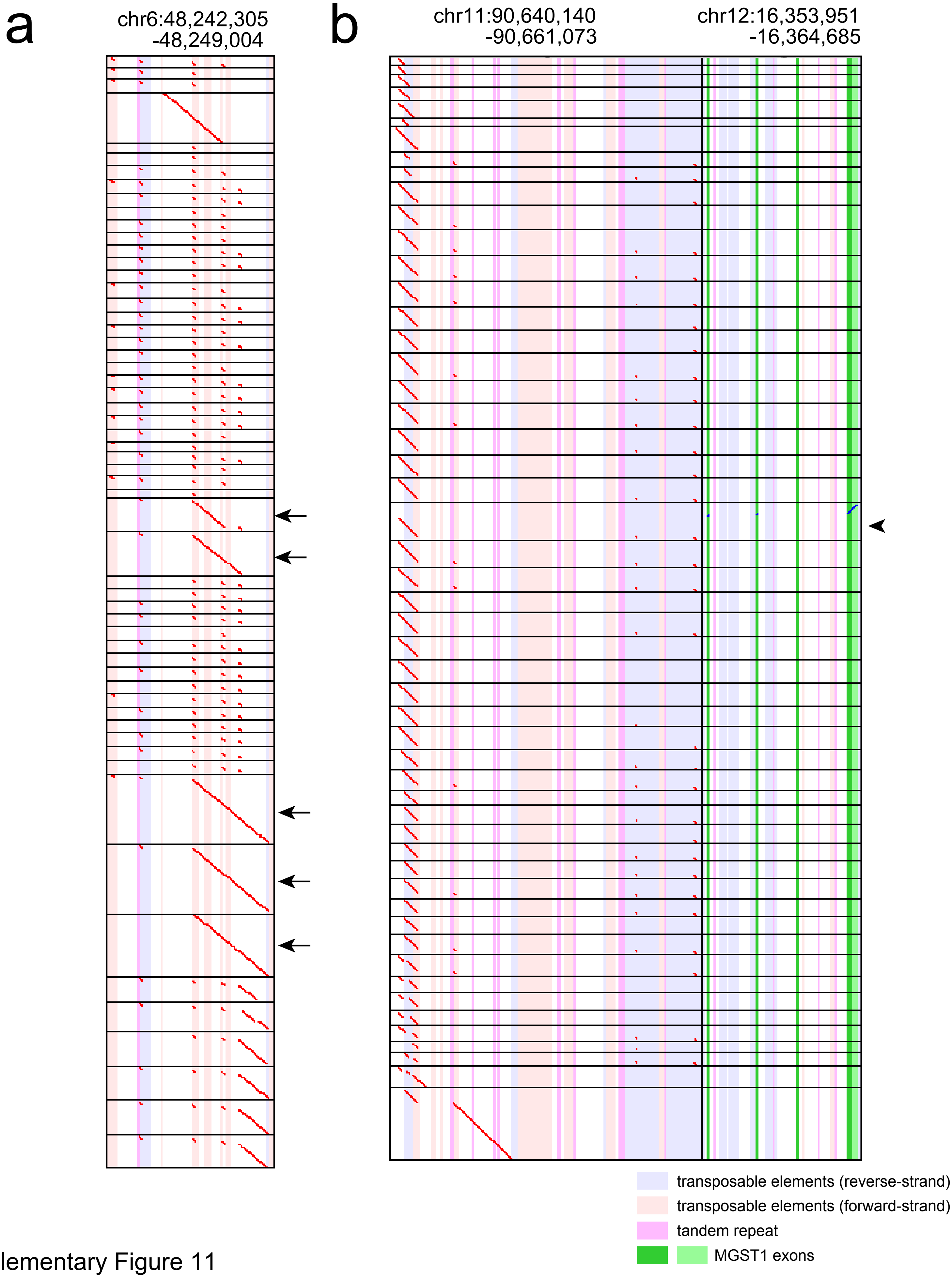


Supplementary Figure 11
(a) Multiple dotplots of actual reads from Figure 3a example transcripts. Note that there are a few reads that were not spliced (arrows). Horizontal black lines separate dotplots of different reads.
(b) The other example of multiple dotplots of spliced transcripts. Arrowhead indicates this transcript fused with exons of a different gene (MGST1).

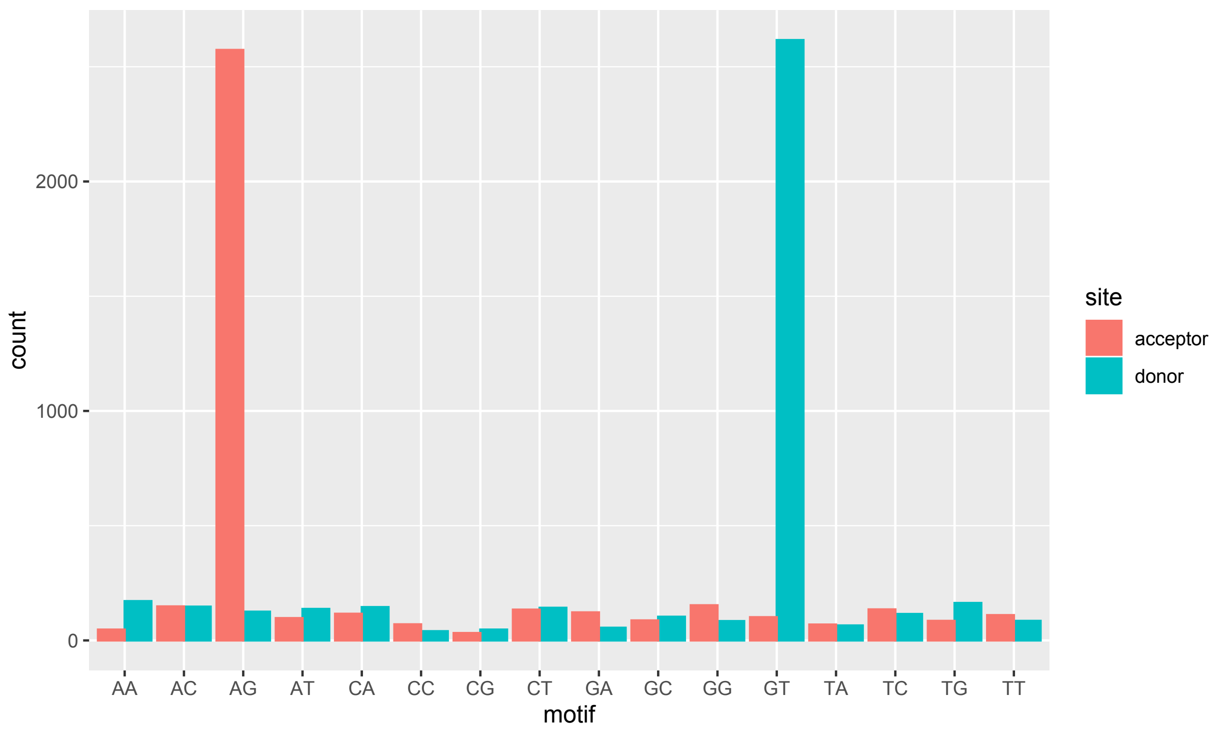


Supplementary Figure 12
Splice site usage predicted from last-rna results. Most of the splicing occurs at AG (acceptor sites) and GT (donor sites).

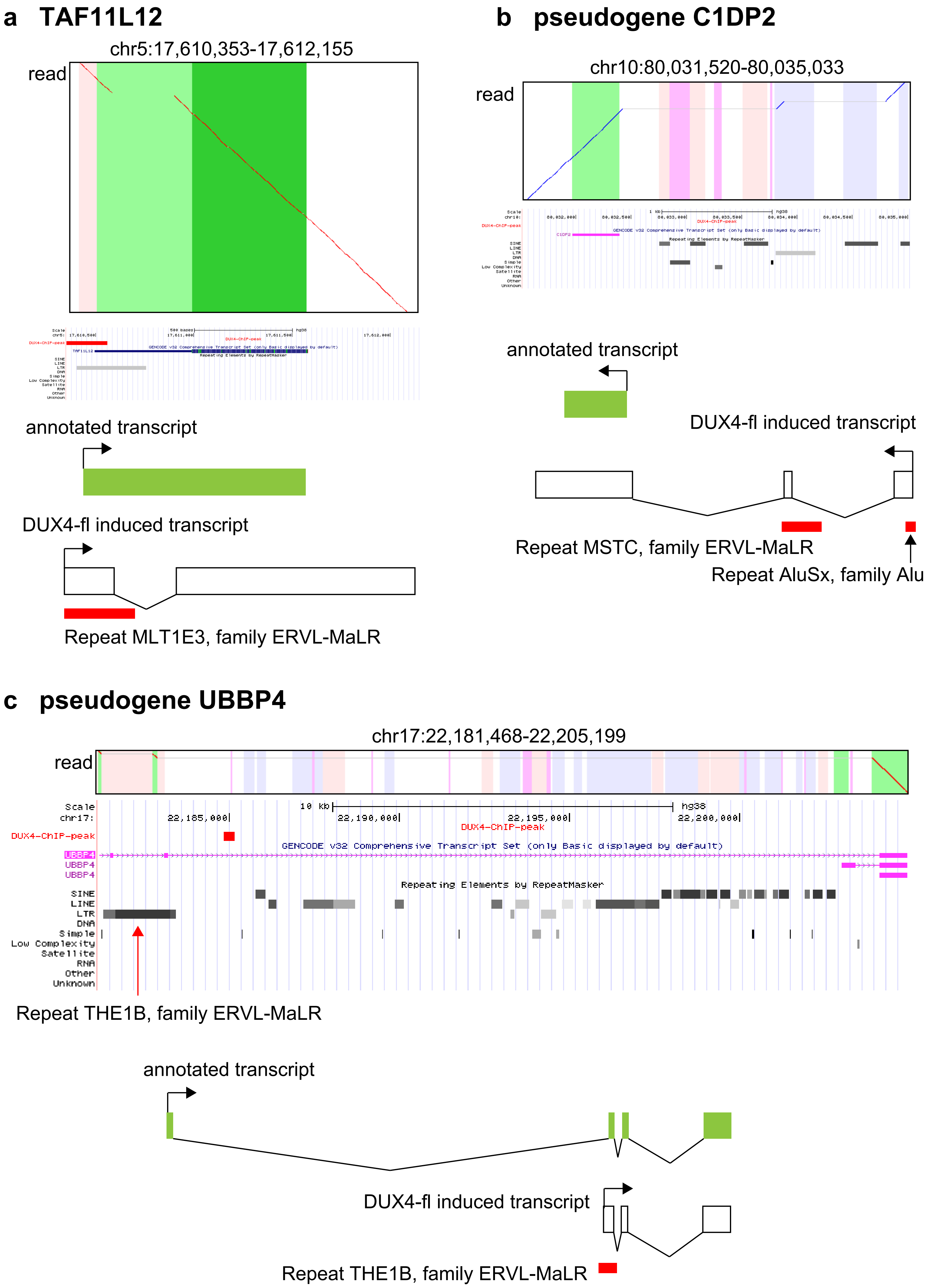
a TAF11L12
chr5:17,610,353-17,612,155
read
annotated transcript
DUX4-fl induced transcript
Repeat MLT1E3, family ERVL-MaLR
b pseudogene C1DP2
chr10:80,031,520-80,035,033
read
annotated transcript
DUX4-fl induced transcript
Repeat MSTC, family ERVL-MaLR
Repeat AluSx, family Alu
c pseudogene UBBP4
chr17:22,181,468-22,205,199
read
Scale
chr17:
10 kb
hg38
22,185,000
22,190,000
22,195,000
22,200,000
DUX4-ChIP-peak
GENCODE v32 Comprehensive Transcript Set (only Basic displayed by default)
UBBP4
UBBP4
UBBP4
Repeating Elements by RepeatMasker
SINE
LINE
LTR
DNA
Simple
Low Complexity
Satellite
RNA
Other
Unknown
Repeat THE1B, family ERVL-MaLR
annotated transcript
DUX4-fl induced transcript
Repeat THE1B, family ERVL-MaLR

**d LINC02229**

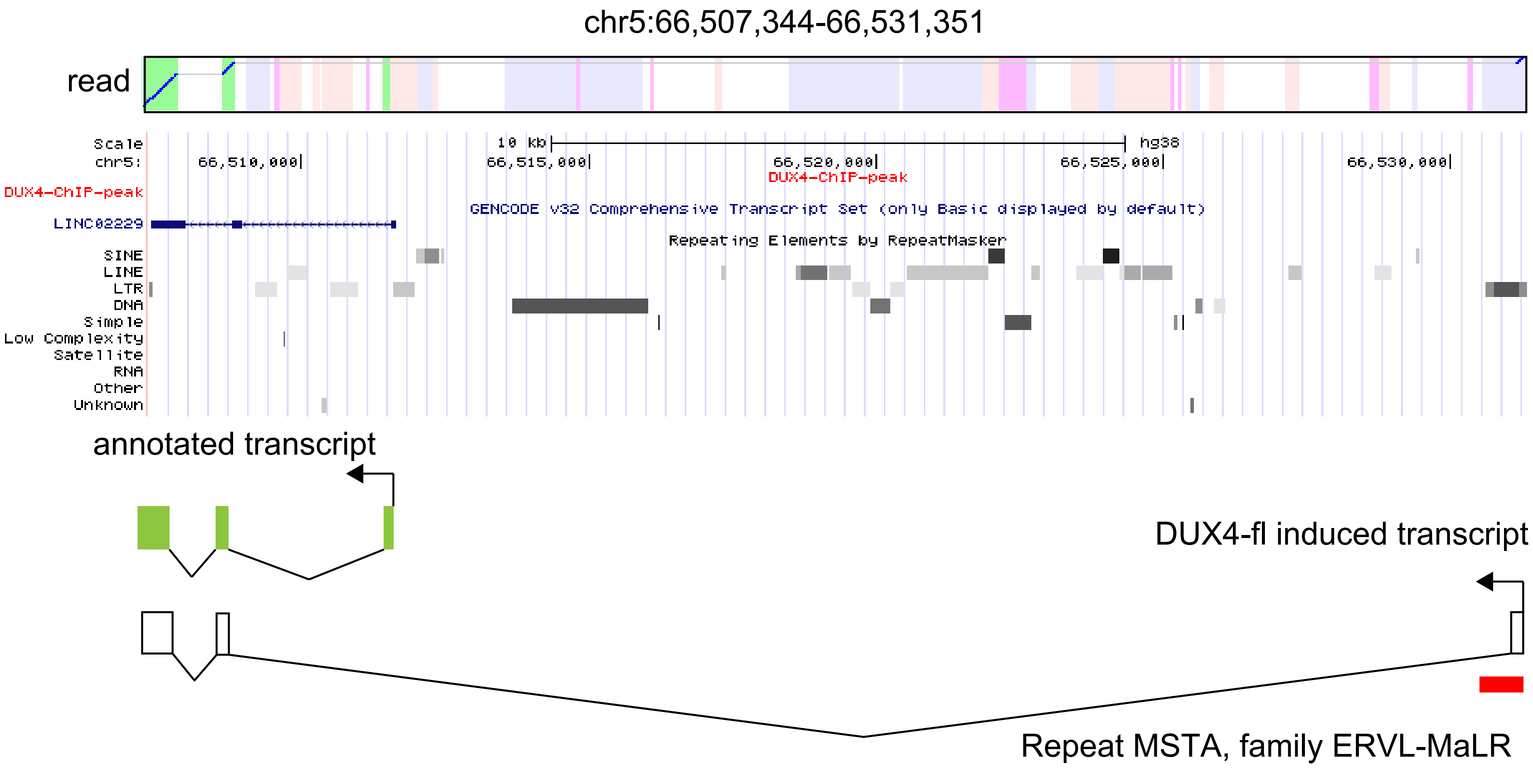


**e LINC02328**

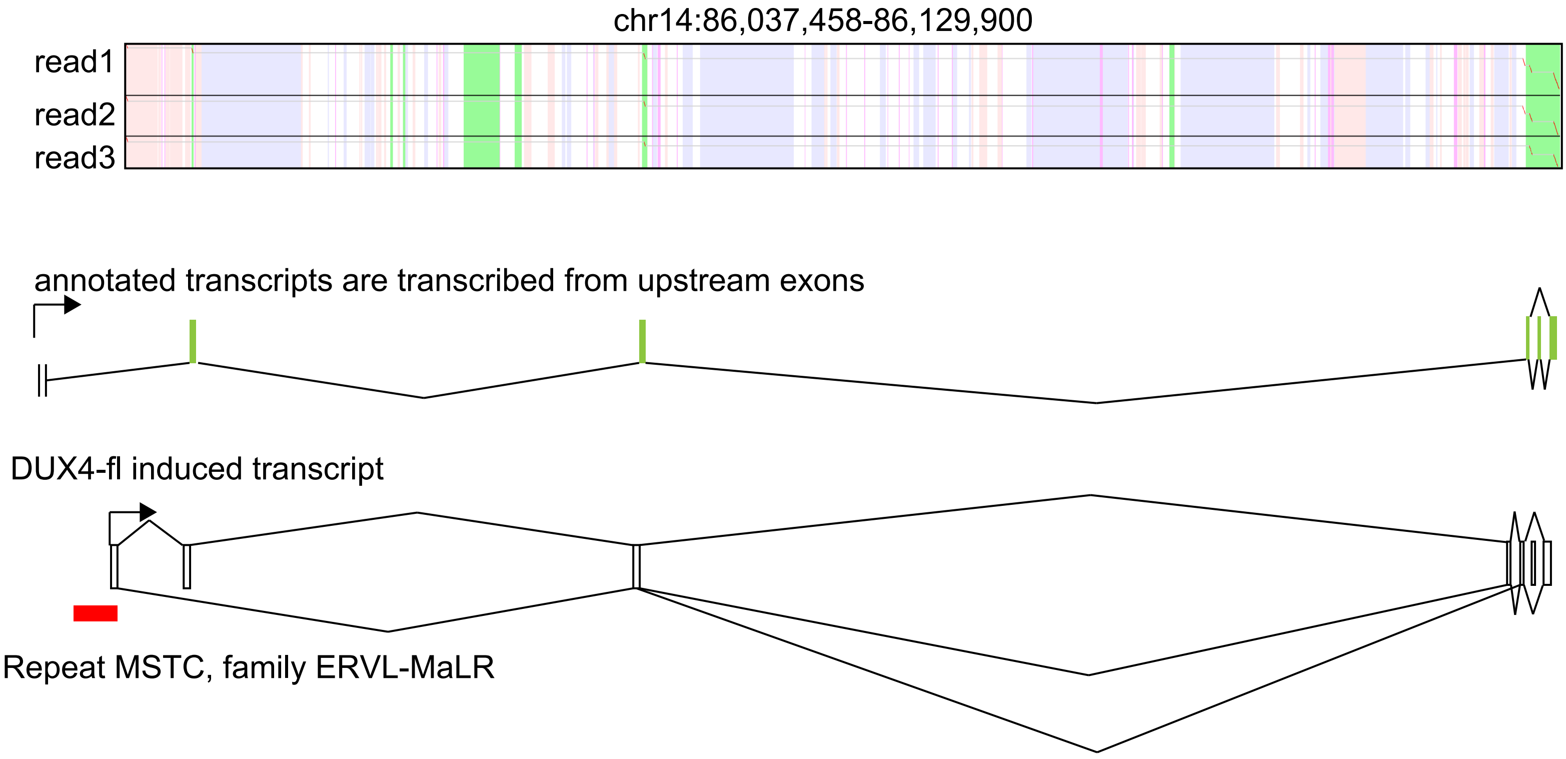

### f RHOBTB1

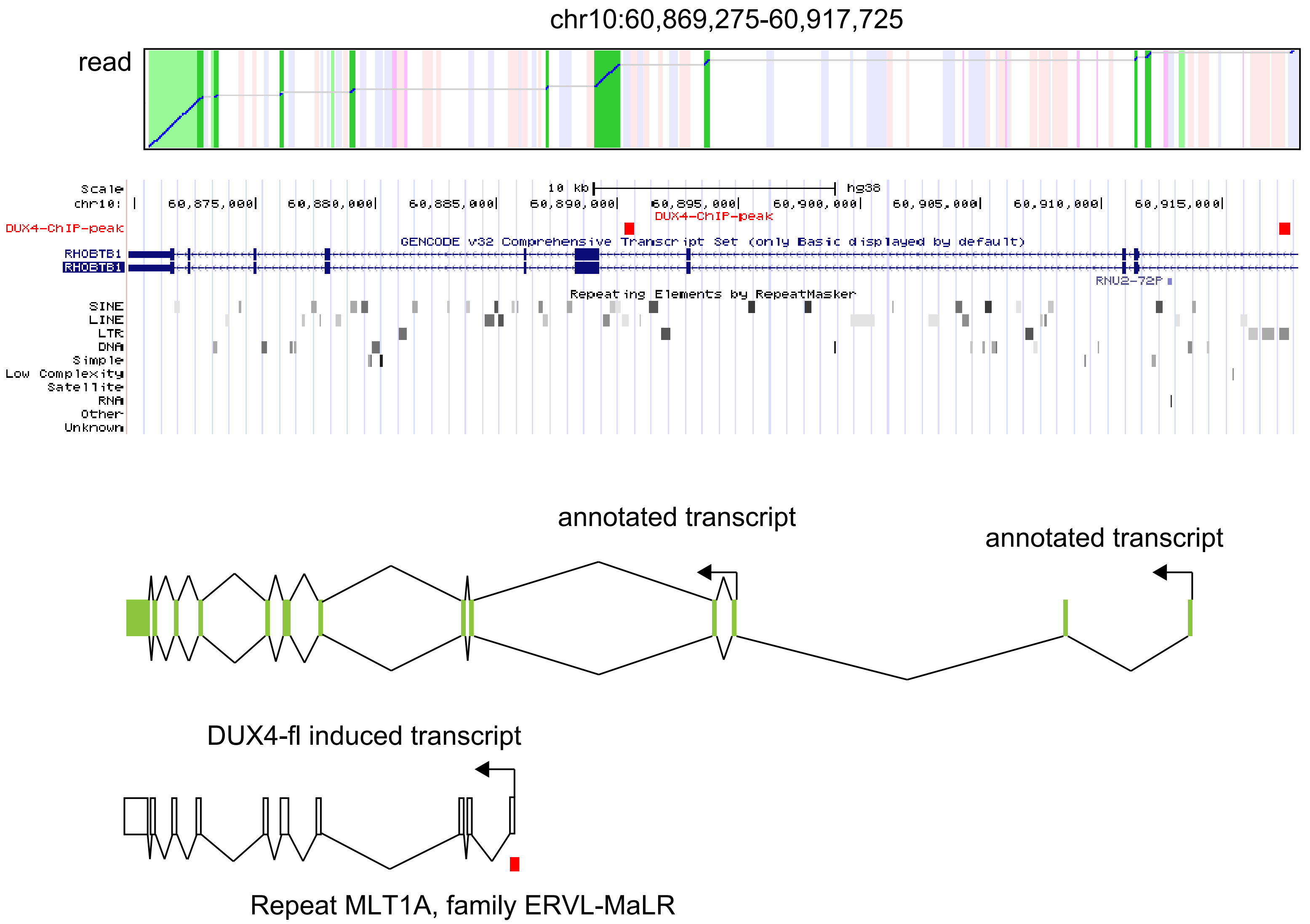

chr10:60,869,275-60,917,725
read
Scale
chr10:
10 kb
hg38
60,875,000
60,880,000
60,885,000
60,890,000
60,895,000
60,900,000
60,905,000
60,910,000
60,915,000
DUX4-ChIP-peak
DUX4-ChIP-peak
GENCODE v32 Comprehensive Transcript Set (only Basic displayed by default)
RHOBTB1
RHOBTB1
RNU2-72P
Repeating Elements by RepeatMasker
SINE
LINE
LTR
DNA
Simple
Low Complexity
Satellite
RNA
Other
Unknown
annotated transcript
annotated transcript
DUX4-fl induced transcript
Repeat MLT1A, family ERVL-MaLR

### g RBM26

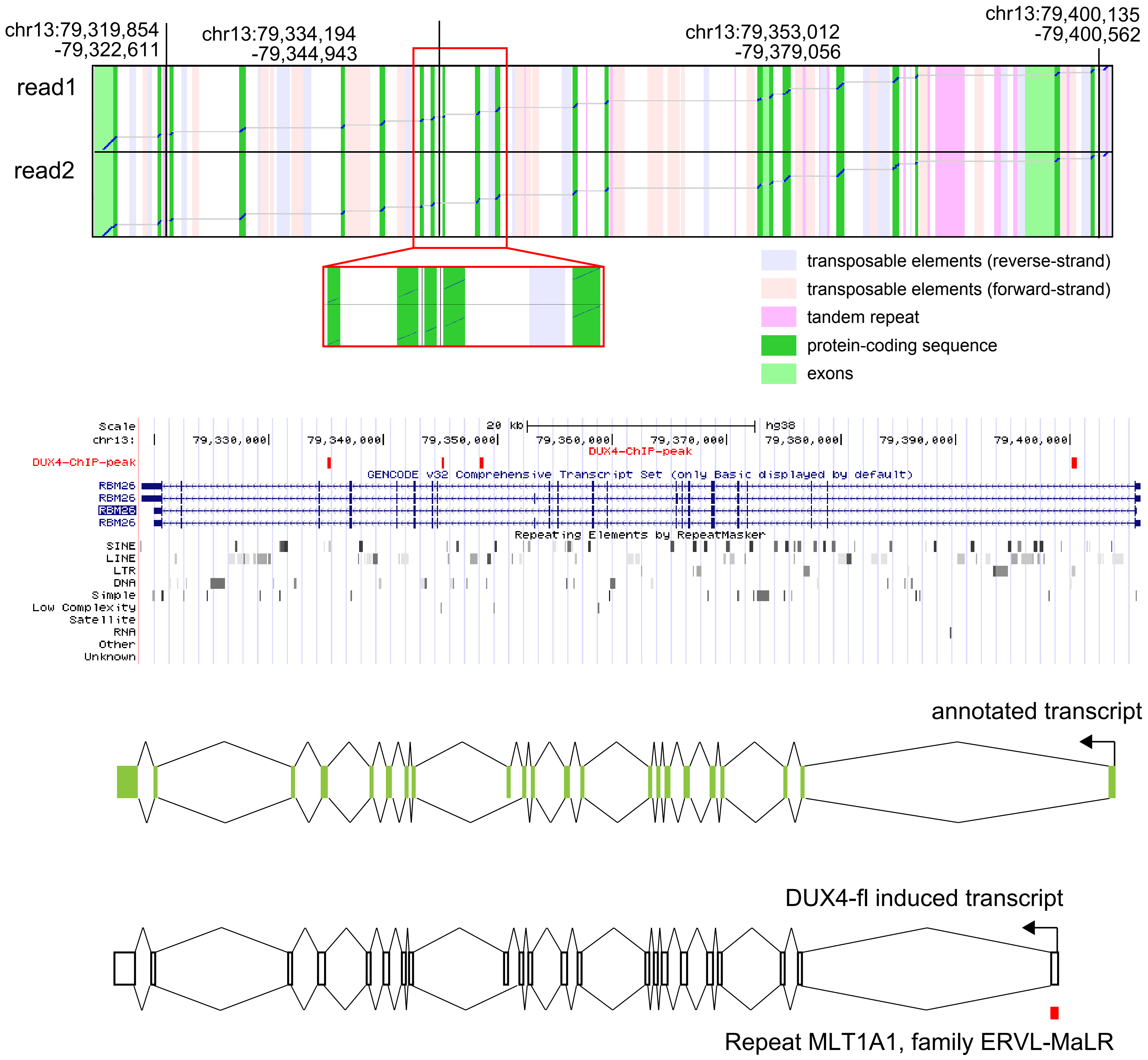


Supplementary Figure 13

(a) TAF11L12 gene was transcribed from its upstream ERVL-MaLR and spliced in the 5' UTR of the gene. The UCSC geneme browser shows a DUX4 ChIP peak, TAF11L12 gene and repeat annotation (Repbase).(b) C1DP2 is a pseudogene (encoded in the minus strand) that was transcribed from its upstream SINE AluSx. (c) Exons 2, 3 and 4 of the UBBP4 pseudogene are transcribed from an ERVL-MaLR located in intron 1 of UBBP4. (d) lncRNA LINC02229 (encoded in the minus strand) is transcribed from its upstream ERVL-MaLR. (e) lncRNA LINC0328 exons are transcribed from an ERVL-MaLR in the intron of the gene, and produce three different splicing isoforms (as shown in read1, 2 and 3). (f) RHOBTB1 (encoded in the minus strand) is transcribed from an ERVL-MaLR in intron 4, creating a new exon 1. Downstream exons were used as annotated.

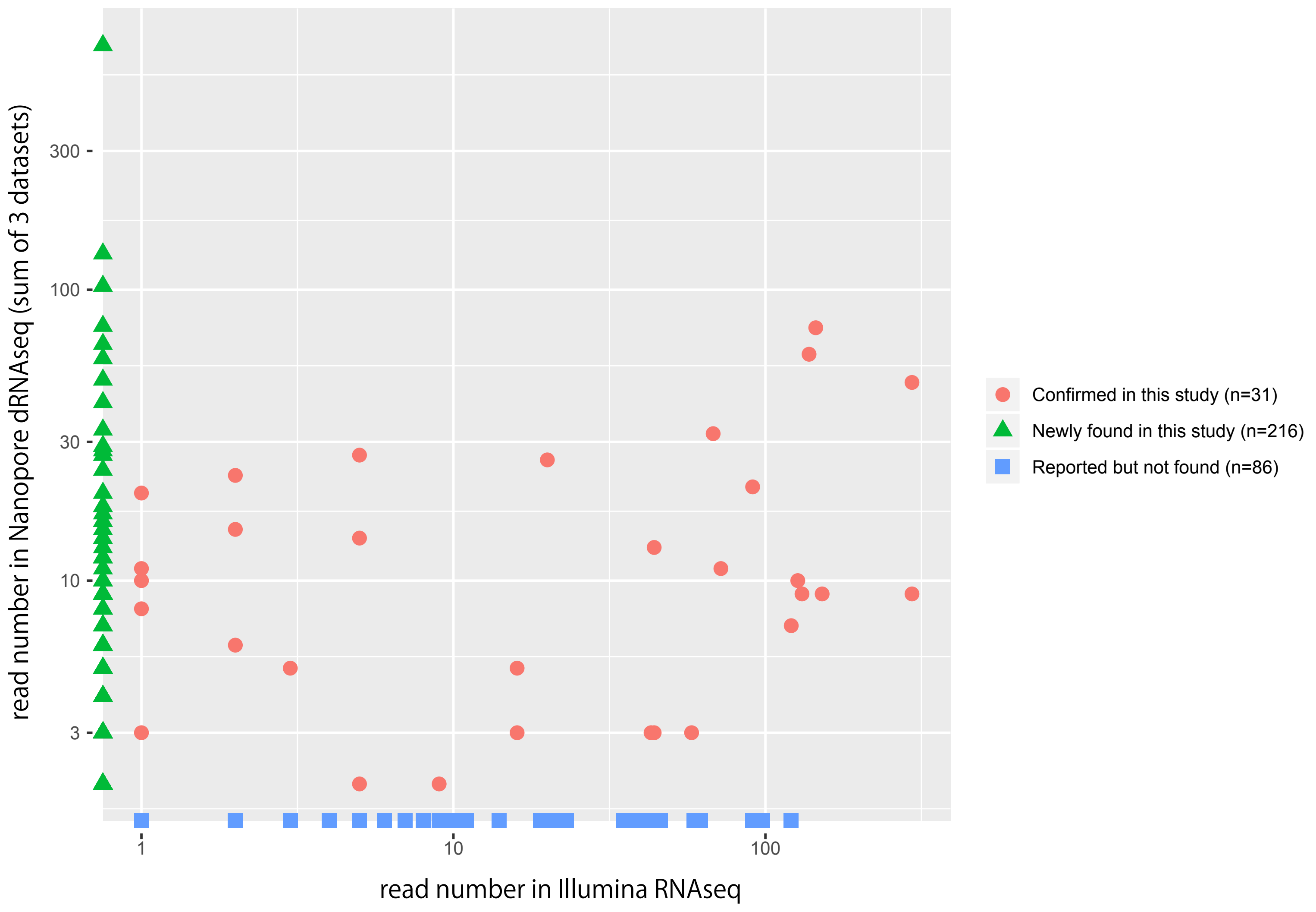


Supplementary Figure 14
The read numbers of DUX4 activated LTR transcripts in dRNA-seq (total of fl-1, fl-2 and fl-3) and previously reported LTR transcripts (14).
